# Molecular, metabolic and functional CD4 T cell paralysis impedes tumor control

**DOI:** 10.1101/2023.04.15.536946

**Authors:** Mengdi Guo, Diala Abd-Rabbo, Bruna Bertol, Madeleine Carew, Sabelo Lukhele, Laura M Snell, Wenxi Xu, Giselle M Boukhaled, Heidi Elsaesser, Marie jo Halaby, Naoto Hirano, Tracy L McGaha, David G Brooks

**Author notes:** Corresponding author: David G. Brooks.

## Abstract

CD4 T cells are important effectors of anti-tumor immunity, yet the regulation of CD4 tumor-specific T (T_TS_) cells during cancer development is still unclear. We demonstrate that CD4 T_TS_ cells are initially primed in the tumor draining lymph node and begin to divide following tumor initiation. Distinct from CD8 T_TS_ cells and previously defined exhaustion programs, CD4 T_TS_ cell proliferation is rapidly frozen in place and differentiation stunted by a functional interplay of T regulatory cells and both intrinsic and extrinsic CTLA4 signaling. Together these mechanisms paralyze CD4 T_TS_ cell differentiation, redirecting metabolic and cytokine production circuits, and reducing CD4 T_TS_ cell accumulation in the tumor. Paralysis is actively maintained throughout cancer progression and CD4 T_TS_ cells rapidly resume proliferation and functional differentiation when both suppressive reactions are alleviated. Strikingly, Treg depletion alone reciprocally induced CD4 T_TS_ cells to themselves become tumor-specific Tregs, whereas CTLA4 blockade alone failed to promote T helper differentiation. Overcoming their paralysis established long-term tumor control, demonstrating a novel immune evasion mechanism that specifically cripples CD4 T_TS_ cells to favor tumor progression.

## INTRODUCTION

Attenuation and deletion of T cell function underlies the inability to control tumor growth and forms the basis for immune-restorative therapies (*1, 2*). Since CD8 T cells are generally the end-point effectors of cancer cell killing, most studies focus on their mechanisms of dysfunction. However, increasing evidence indicates the importance of CD4 T helper (Th) cells in controlling tumor growth and increasing efficacy of immunotherapies (*3–6*), suggesting a need to understand their dysfunction during cancer development. Interestingly, CD4 T_TS_ cells and CD8 T_TS_ cells possess different sensitivity towards immune checkpoint blockades (*7*), and as a result, likely are mediated by different regulatory mechanisms. Following activation, naïve CD4 T cells differentiate into specific subsets, based on signals from the antigenic environment and interactions with antigen presenting cells (APCs). The CD4 Th response is guided to control individual types of immune responses, with misdirection potentially leading to ineffective disease control (*8–11*). Therefore, it is important to understand the activation, differentiation and regulation of CD4 T_TS_ cells in cancer.

Interestingly, a growing number of studies in humans indicate that only a fraction of tumor-infiltrating T lymphocytes (TILs) are in fact tumor-specific, and although generally limited to analysis of CD8 T cells, much of what may be considered tumor-specific T cells based on phenotypes (e.g., inhibitory or stimulatory receptors) are instead specific to other antigens, such as virus-specific T cells (*12–15*). The frequency of tumor-reactive CD4 and CD8 TILs can be increased by immunotherapy or *ex vivo* expansion for adoptive cell therapy (*4, 16*), suggesting immune regulatory mechanisms that limit the effective accumulation of T_TS_ cells in the TME. However, so far studies using mouse tumor models to understand anti-tumor CD4 T cell function have primarily focused on studying bulk CD4 TILs with unknown antigen specificities (*7*), transferring CD4 T cells to lymphopenic recipients with pre-established tumors (*17, 18*), or transferring *in vitro* activated CD4 T cells as adoptive cell therapy (ACT). Thus, how naïve CD4 T_TS_ cells are initially activated and then differentiate in response to tumor progression is not well understood and will be important for the development of strategies to invoke CD4 T_TS_ cellsto control cancer growth.

Herein, we demonstrate that CD4 T_TS_ cells in the draining lymph node (dLN) were initially primed following tumor initiation but were then rapidly frozen into a long-term paralyzed state distinct from effector or exhaustion programming, that impeded tumor control. The persistent paralysis state limited the tumor homing and accumulation of CD4 T_TS_ cells, resulting in only a small presence of CD4 T_TS_ cells in the tumor compared to CD8 T_TS_ cells. Molecular and cellular interrogation of the paralyzed state identified unique programs potentiating CD4 T_TS_ cell paralysis, including abrogated Th fate commitment, failure to produce IFNγ, an altered metabolic state limiting aerobic glycolysis while pushing toward oxidative phosphorylation, reduced mitochondrial fitness, and increased pathways of cellular stress and cell death. Unlike the well-defined states of cellular exhaustion, the paralysis was not due to chronic antigenic stimulation or PD1:PDL1 interactions, but instead was mediated by a coordination of Treg cells and CTLA4 signalling that continually restricted CD4 T_TS_ cell activation throughout cancer progression. In contrast, CD8 T_TS_ cells did not exhibit the proliferative paralysis or tumor homing inhibition. Overcoming the paralysis enabled robust CD4 T_TS_ cell tumor infiltration and enhanced tumor control, thus defining a novel CD4 T cell directed mechanism of tumor-driven immune escape that preferentially inhibits CD4 T_TS_ cells.

## RESULTS

### CD4 T cells are critical for anti-tumor immune responses and their amplification can enable long-term tumor control

To understand how the CD4 T cell response is regulated in response to tumors, we identified murine tumor models wherein CD4 T cells contributed to long-term control. While the growth of B16-F10 (melanoma) was not affected by the absence of CD4 T cells, both PyMT (orthotopic breast tumor) and MC38 (colon adenocarcinoma) exhibited accelerated growth in CD4 knockout (KO) mice compared to wildtype (WT) mice (Figure 1A, S1A). This accelerated growth in CD4 KO mice was counter to what would be expected from the loss of Tregs alone; and indeed, early Treg depletion did enhance MC38 and PyMT tumor control (Figure S1B), indicating that the absence of conventional CD4 T cells in the CD4KO mice outweighed the beneficial effect of Treg depletion. Based on its requirement for CD4 T cells and prolonged growth time (∼40-60 days to reach endpoint) allowing for the establishment of a chronic tumor: immune interaction, we primarily focused on the PyMT model. Results were then confirmed using the MC38 tumor model.

**Figure 1.**
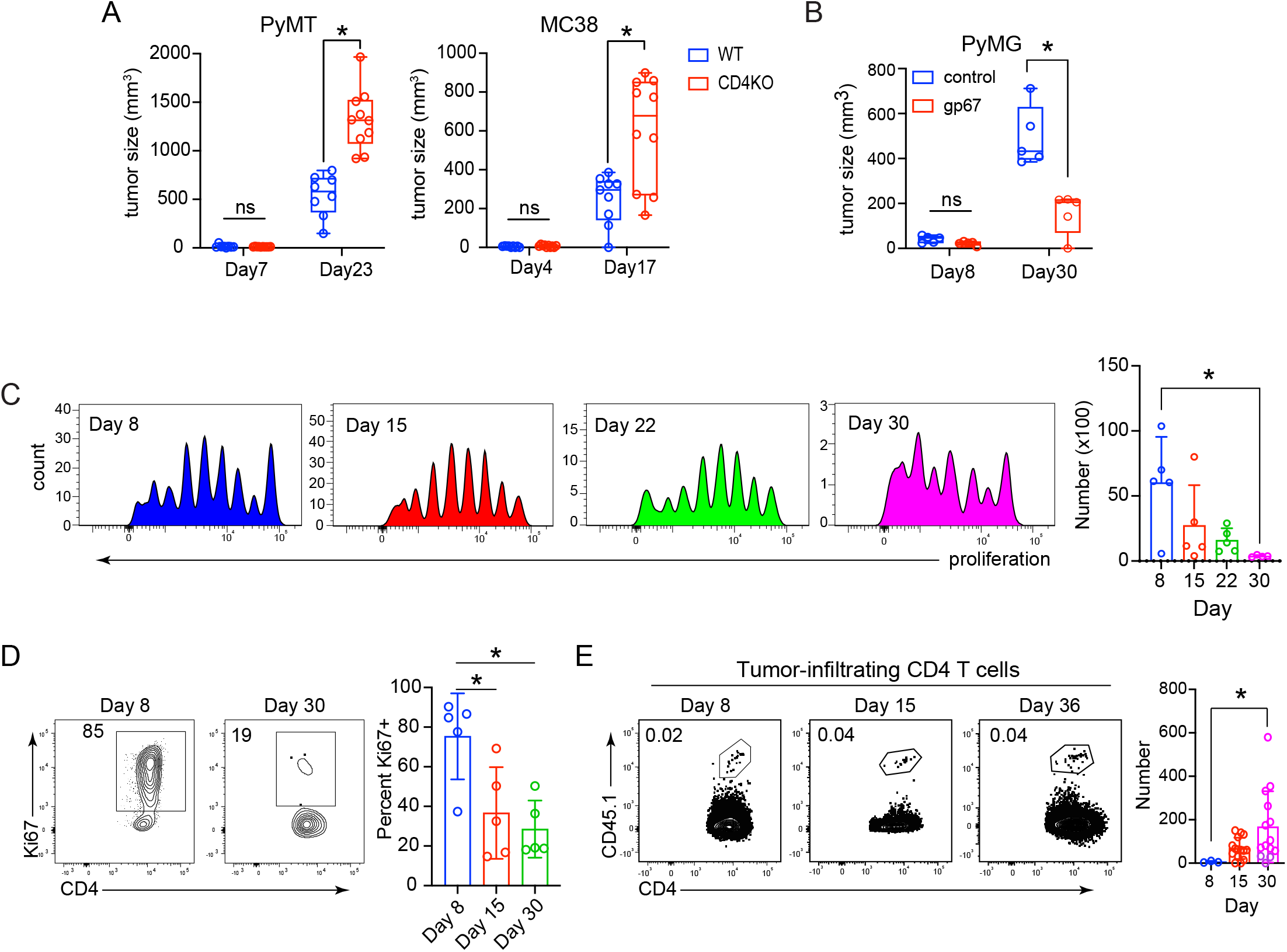
Sub-optimal help and rapid proliferative paralysis of CD4 T_TS_ cells. (A) Tumor sizes of PyMT and MC38 tumor cells in WT and CD4 knockout (KO) mice at indicated time points. (B) Mice received naïve CD4 SMARTA T_TS_ cells followed by implantation of PyMG tumors. On the same day of tumor implantation, mice received unlabeled, or GP_61-80_ peptide labeled β2m-/- bmDC. Bar graph shows the tumor size at the indicated day. (C-E) TAGIT proliferation dye labelled naïve SMARTA cells were transferred into WT mice followed by PyMG injection. (C) Histograms show TAGIT dilution on the indicated day. The bar graph represents the total number of CD4 SMARTA T_TS_ cells in the dLN at the indicated day after PyMG administration. (D) Expression (flow plot) and the percentage of Ki67+ out of total CD4 SMARTA T_TS_ cells (gated on SMARTA cells) in the dLN. (E) Representative percentage (flow plots) and number (bar graph) of CD4 SMARTA T_TS_ cells in the tumor at each time point. Data represents 3 independent experiments with at least 5 mice per group. Error bars indicate standard deviation (SD). For comparison of tumor growth kinetics, significance is determined by two-way ANOVA; for other comparisons, significance is determined by Mann-Whitney U test. *, p<0.05.

It is difficult to distinguish tumor-specific from “bystander” CD4 T cells within the tumor and dLN based on phenotypic proteins. To identify CD4 T_TS_ cells *in vivo*, we created PyMT and MC38 cell lines expressing the LCMV-glycoprotein (GP)_1-100_ sequence. These new lines (PyMG and MC38GP) express the LCMV-GP_61-80_ epitope and the LCMV-GP_33-41_ epitope as model tumor antigens to allow *in vivo* identification of endogenous CD4 and CD8 T_TS_ cells as well as the use of their TCR transgenic counterparts (SMARTAs for CD4 T cells and P14s for CD8 T cells) that are frequently used to reliably understand T cell dysfunction in chronic viral infections and cancers (*6, 19, 20*). The new tumor lines develop into lethal tumors in regular C57BL/6 mice, indicating that insertion of the LCMV-derived sequences did not significantly enhance the immunogenicity of these tumors in the single-cell clones used herein (Figure S1C). We first determined whether enhancing CD4 T_TS_ cell help would increase tumor control. For this, bone marrow-derived DC (bmDC) were generated from β2microglobulin-/22mice (β2m-/-, lacking MHC I expression and incapable of stimulating CD8 T cells) and pulsed with MHC II-restricted LCMV-GP_61-80_ peptide or left unlabeled. In this way, only the GP_61-80_ specific CD4 T cells are activated by the bmDC. Transfer of the GP_61-80_ peptide labeled β2m-/- bmDC alone decreased PyMG tumor growth (Figure S1D), and tumor control was further enhanced by transferring naïve TCR transgenic GP_61-80_ specific CD4 T cells (i.e., SMARTA cells) to increase the initial precursor frequency of CD4 T_TS_ cells (Figure 1B). The transfer of GP_61-80_-labeled β2m-/- bmDCs also increased and sustained tumor-specific CD4 SMARTA T cell tumor infiltration (Figure S1E). In contrast, tumor infiltration by CD4 T_TS_ cells was almost entirely absent in the mice receiving unlabeled bmDC (Figure S1E). Further, transferring GP_61-80_-labeled β2m -/- bmDCs 21 days after tumor initiation induced better tumor control (Figure S1F). Together these observations with bmDCs and CD4KO mice indicate that while exerting a beneficial effect, without therapeutic activation the naturally-induced CD4 T_TS_ cell response remains sub-optimal.

### CD4 T_TS_ cell proliferation paralysis is rapidly induced in the draining lymph node

To understand the basis for the sub-optimal CD4 T cell help, we adoptively transferred naïve CD4 SMARTA T_TS_ cells prior to PyMG implantation. By day 8 after PyMG implantation (when the tumor was barely palpable, < 50 mm^3^), about 40% of the CD4 SMARTA T_TS_ cells in the dLN had divided (Figure 1C, Figure S2A). Further, the proliferated cells upregulated activation-induced proteins CD44, CD86, and Ki67, indicating cell cycling (Figure 1D, S2B). Despite of the division and cell activation within the dLN, only a small portion of fully-divided (i.e. completely diluted cell proliferation dye) CD4 T_TS_ cells infiltrated the tumor by day 8, and the amounts did not substantially increase with tumor progression (Figure 1E, S2C).

The initial activation failed to sustain cell division in the dLN. Instead the CD4 T_TS_ cells became “paralyzed” in their division cycle, with a progressive loss of actively cycling Ki67+ cells and minimal subsequent cell division through tumor progression (Figure 1C, 1D). Tumor progression was accompanied by a numerical decrease in the number of CD4 T_TS_ cells in the dLN, with the reduction similarly observed in each cell division peak (Figure 1C, S2D). Furthermore, the loss of CD4 T_TS_ cells in the dLN was not accompanied by a comparable appearance in the tumor (Figure 1E), suggesting that the decreased number was due to an attrition of CD4 T_TS_ cells as tumor growth progressed. A similar proliferative paralysis was observed for tumor-specific CD4 SMARTA T cells responding to MC38GP tumors, and for OT-II cells responding to ovalbumin-expressing MC38 tumors (Figure S2E), demonstrating that the proliferative paralysis occurs in multiple tumor models and with different TCR specificities. In contrast, CD8 T_TS_ cells continued to proliferate, were numerically sustained in the dLN, and progressively infiltrated the tumor (Figure S2F, S2G), indicating that the proliferative paralysis was specific to CD4 T cells. Importantly, the proliferative paralysis was not due to loss of GP_61-80_ expression nor a progressive failure of APCs to present antigen *in vivo* since naïve CD4 SMARTA T cells transferred 21 days after tumor-initiation were activated and proliferated (Figure S2H). However, these late-transferred CD4 T_TS_ cells also experienced a proliferative paralysis (Figure S2H), indicating the continual maintenance of mechanisms to initially allow priming, but then paralyze the CD4 T_TS_ cells. Importantly, the priming at 21 days after tumor initiation shows that neither the cell transfer nor provision of dead cells at the time of tumor transplant are responsible for the paralysis, but instead that this is a physiologic response of CD4 T cells to tumor priming and growth.

### CD4 T_TS_ cell differentiation is stunted, providing restricted help through tumor progression

In addition to their proliferative paralysis, CD4 T_TS_ cells in the dLN at the early stage of tumor (Day8) failed to express most Th-defining transcription factors, including Tbet (Th1), RORgT (Th17) or FoxP3 (Treg) (Figure 2A, S3A). Conversely, the majority of the CD4 T_TS_ cells expressed and maintained high levels Bcl6 and TCF1 (Figure 2A). Virus-specific CD4 T cells from chronic LCMV-Clone 13 (Cl13) infection are shown for comparison to identify Th1 (Tbet+) and Tfh (Bcl6+) and the expression of these lineage defining proteins (Figure 2A). In comparison, Bcl6 was not expressed in the naïve CD4 T cells, while TCF1 was continuously expressed at high levels in CD4 T_TS_ cells (Figure 2A, S3B), indicating that the CD4 T_TS_ cells are being activated, but not sufficiently to down-regulate TCF1 expression from naïve cell levels. This diminished activation level is further supported by the low expression of the TCR-induced costimulatory molecule CD27 (*21*) (Figure 2A). Increased Bcl6 and TCF1 are associated with Tfh differentiation, however, the CD4 T_TS_ cells had much lower expression of other Tfh lineage-defining and functional proteins, including CXCR5 or ICOS compared to the virus-specific Th1 and Tfh that are generated during chronic LCMV infection (Figure 2B), suggesting that these cells are not Tfh-committed. Further, the CD4 T_TS_ cells produced high levels of IL2 and TNFα in response to antigen, but minimal IFNγ or IL10 (Figure 2C, S3C, S3D), indicating restricted, yet specific, cytokine producing potential through cancer progression. By 30 days after PyMG initiation, 10- 40% of the CD4 T_TS_ cells in the dLN had upregulated FoxP3, with most of the FoxP3-expressing cells having proliferated extensively (Figure 2D). Within the tumor, despite low amounts of CD4 T_TS_ cells, the majority expressed FoxP3 at day 30 but not day 8 (Figure 2D), suggesting that tumor-specific CD4 Treg cells preferentially and progressively accumulate in the tumor.

**Figure 2.**
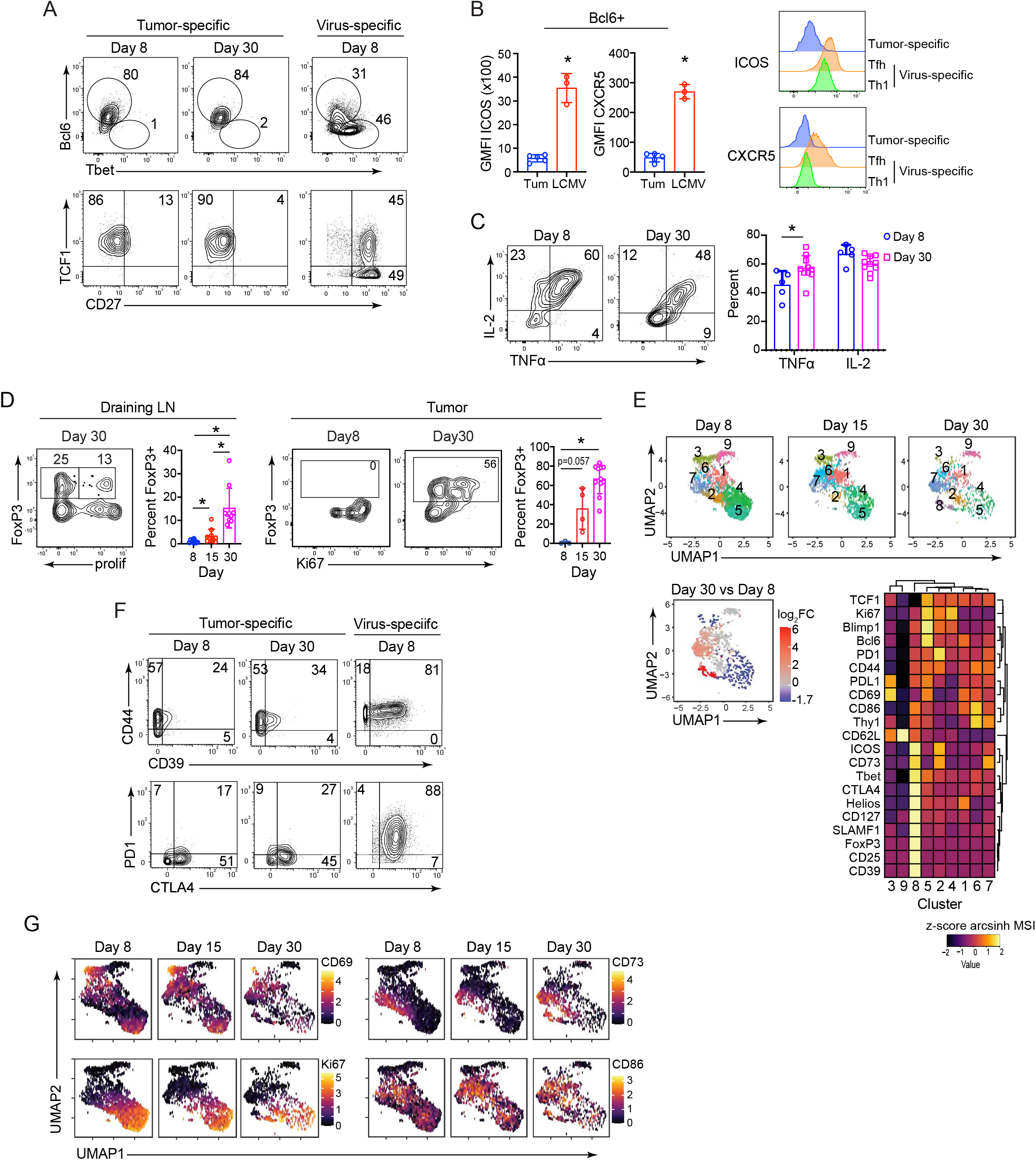
Proliferative paralysis and functional restriction of CD4 T_TS_ cells. (A) Representative expression of the indicated protein CD4 SMARTA T_TS_ cells (in the dLN) on the indicated day. Virus-specific CD4 SMARTA T cells from chronic LCMV infection are shown for comparison. (B) Geometric mean fluorescence intensity (GMFI) of ICOS and CXCR5 by Bcl6+ tumor-specific or virus- specific CD4 SMARTA Tfh cells (bar graphs). Histograms show representative ICOS and CXCR5 expression by dLN tumor-specific SMARTA cells, and by virus-specific CD4 Th1 and Tfh cells. (C) Expression and quantification of TNFα and IL2 producing CD4 SMARTA T cells in dLN after ex vivo LCMV-GP_61-80_ peptide stimulation. (D) (Left) FoxP3 versus TAGIT expression by CD4 SMARTA T_TS_ cells on day 30 in the dLN. The bar graph is gated on CD4 SMARTA T_TS_ cells and shows the percentage of FoxP3+ cells in the dLN. (Right) Plots show the cycling (Ki67+) FoxP3+ CD4 SMARTA T_TS_ cells in the PyMG tumor. The bar graphs show the percentage of FoxP3+ SMARTA T_TS_ cells in the tumor. (E) (Top) UMAP embeddings of CyTOF data of CD4 SMARTA T_TS_ cells in the dLN at day 8, 15 and 30 after PyMG initiation. Cells are colored by Phenograph clustering. Data is concatenated from 5-7 mice per group. (Bottom) UMAP plot shows the clusters that are increased (red) or decreased (blue) in abundance from day 8 to day 30. The bar graph indicates the log2 fold change of each cluster and the associated significance (calculated using diffcyt (*69*), * adjusted p value <0.05). Heatmap depicts the normalized z- scores of the arcsinh-transformed median signal intensity (MSI) of the indicated protein in each cluster. (F) Expression of the indicated proteins by CD4 SMARTA T_TS_ cells in the dLN. Virus-specific CD4 SMARTA T cells from chronic LCMV infection are shown for comparison. (G) UMAPs show the arcsinh-transformed single-cell expression of the indicated proteins by CD4 SMARTA T_TS_ cells in dLN at each time point. For flow cytometry, data represents 3 independent experiments with at least 5 mice per group. Error bars indicate SD. Significance is determined by Mann-Whitney U test. *, p<0.05. CyTOF data are representative of 2 experiments with at least 3 mice per group.

We next performed mass cytometry (CyTOF) analyses of CD4 T_TS_ cells in the dLN throughout PyMG progression. Phenograph-based clustering of the CyTOF data resolved nine clusters that could be broadly grouped into three major categories: (1) naïve-like cells expressing high CD62L, in combination with low activation-induced proteins [cluster (c) 9]; (2) activated populations (c1-7); and (3) induced Treg (iTreg) cells (c8) (Figure 2E). Despite the ongoing antigen presentation, the naïve (c9) and all the effector populations of CD4 T_TS_ cells (c1-7) remained relatively stable with limited changes in their expression of activation, inhibitory, or Th-lineage defining transcription factors during tumor progression (Figure 2E, S3E, S3F). The exception was the appearance between days 15 and 30 of a tumor-specific FoxP3+ Helios+ iTreg population (c8) that expressed proteins associated with Treg function (e.g., CD25, CD73, CD39, CTLA4, CD39), and in general had the highest activation level among all tumor-specific CD4 T cell clusters (Figure 2E, S3E). Interestingly, the iTregs also expressed Th1-lineage defining proteins, including Tbet and SLAMF1 (Figure 2E, S3G), suggesting tumor-specific iTregs develop Th1-like features that could further limit antigen-specific CD4 and CD8 T cells (*9*).

To gauge the expression level of activation-induced proteins on CD4 SMARTA T_TS_ cells, we compared them to the same CD4 SMARTA T cells activated during chronic LCMV-Cl13 infection. Unlike the virus-specific SMARTA T cells that expressed high levels of activation-induced and inhibitory proteins, the CD4 T_TS_ cells expressed much lower levels of these with many not expressing them at all, including CD39 or PD1 (Figure 2F). Yet, a large proportion of the CD4 T_TS_ cells expressed CTLA4 (albeit less than in the chronic virus infection) (Figure 2F) suggesting preferential expression of T cell inhibitory receptors. Deeper analysis indicated that the activated populations could be further categorized based on the expression of four dominating protein expression profiles: (1) CD69-hi, CD62L-int, recently-activated cells (c3); (2) Ki67+ cycling cells (c4 and c5); (3) CD73-hi populations (c2 and c7); and (4) non-cycling CD86-hi populations (mostly c1 and c6, but also have some overlap with the CD73 and the Ki67 expressing populations) (Figure 2G). Consistent with progressive loss of Ki67 expressing cells, c4 decreased in abundance rapidly (between day 8-15) while c5, which had upregulated Bcl6 and other activation induced proteins, persisted longer (decreasing between day 15 to 30) (Figure S3E). While the CD86-hi c1 and c6 remained stable within the CD4 T_TS_ cell population, the frequency of CD73-hi clusters increased during the late stage of tumor progression (Figure 2G, S3E). Counter-intuitively, Bcl6 and Blimp1 [transcription factors that generally repress each other (*22*)], were co-expressed by the Ki67+ cells (Figure S3H), further highlighting the uncommitted differentiation states in the activated CD4 T_TS_ cells.

### Altered metabolic state is associated with CD4 T_TS_ cell paralysis state

Metabolic reprogramming from oxidative phosphorylation to aerobic glycolysis after activation (the Warburg effect) is critical for T cell proliferation, differentiation, and function (*23, 24*). To probe single-cell changes in CD4 T_TS_ cell metabolism, we adapted a metabolism CyTOF panel from human to mouse cells (*25*). We compared naïve CD4 SMARTA T cells and CD4 SMARTA T cells on day 8 after PyMG initiation, as well as CD4 Th1 and Tfh SMARTA cells from day 8 after acute LCMV-Armstrong (Arm) infection. We used the virus-specific CD4 T cell response to acute LCMV-Arm infection because it is a gold-standard to define bona fide CD4 Th1 and Tfh differentiation (*26*). In this way, the same CD4 SMARTA T cells were analyzed in various conditions, thus allowing direct comparison of T cell metabolic profile. Compared to LCMV-specific Th1 and Tfh cells, key proteins involved in glucose uptake, GLUT1, and the glycolysis pathway, hexokinase 2 (HK2), and GAPDH, had decreased expression in CD4 T_TS_ cells (Figure 3A-C, S4A), suggesting an inhibited switching to aerobic glycolysis for energy production. Interestingly, GAPDH, a key enzyme involved in glycolysis and an important transcription regulator of T cell function(*27*), was reduced even compared to naïve T cells (Figure 3A, 3B, S4A). PFKFB4, the bi-functional kinase/phosphatase that indirectly affects glycolysis by regulating the concentration of glycolytic byproduct fructose-2,6-biphosphate, was up-regulated in tumor-activated CD4 T cells (Figure 3A-C, S4A). The upregulation of PFKFB4 has been reported to be induced by hypoxia via HIFa to promote further uptake of glucose in many cancer cells and promote their survival (*28, 29*). Though the function of PFKFB4 in T cell is still not clear, its upregulation in CD4 T_TS_ cells as compared to virus-specific CD4 Th1 and Tfh cells could serve as a compensatory response to the reduced glycolysis pathway. Thus, CD4 T_TS_ cells fail to effectively transition to aerobic glycolysis, which could hinder their proliferation and survival.

**Figure 3.**
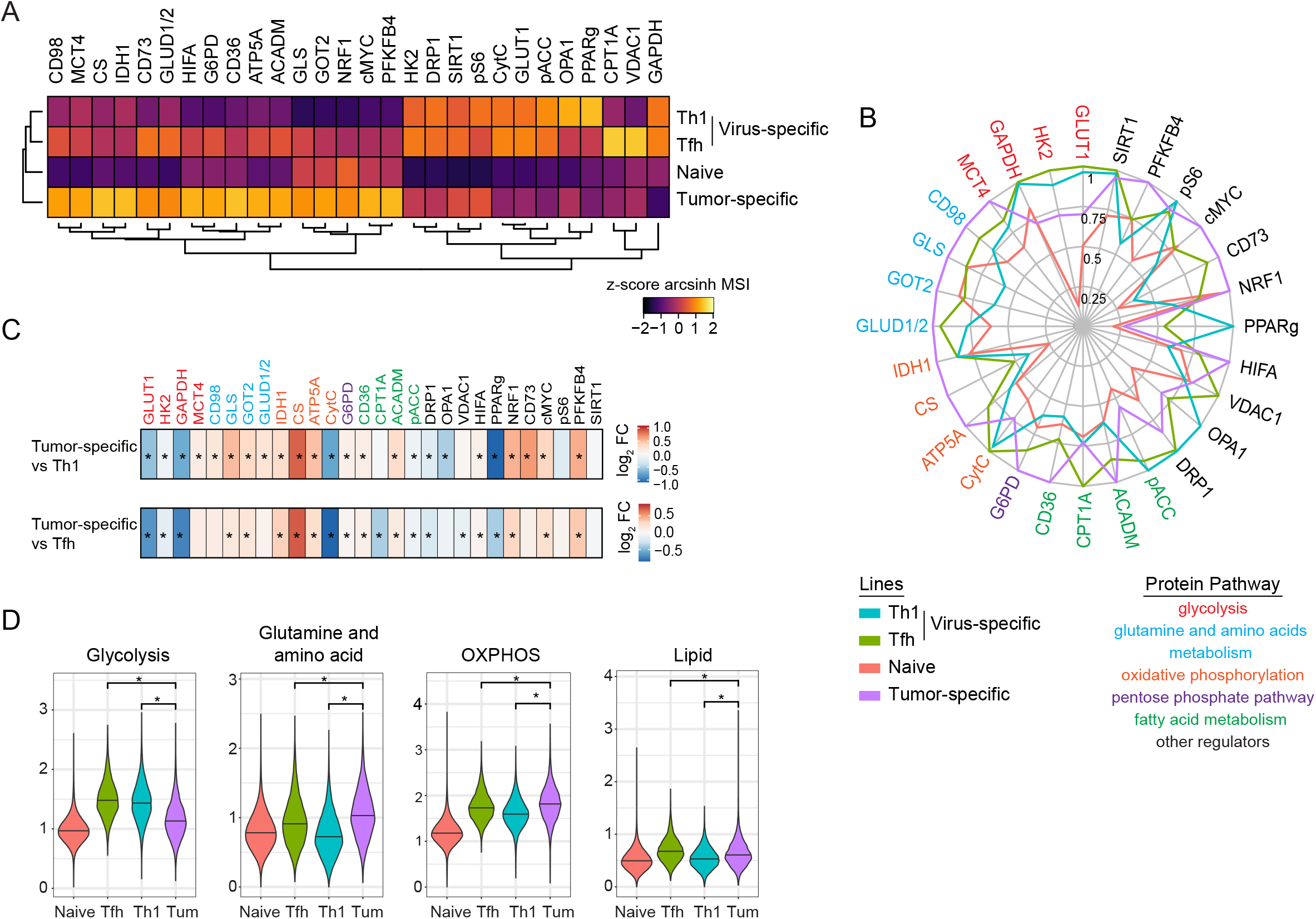
CyTOF analyses of CD4 T_TS_ cell metabolic pathways. CD4 SMARTA T cells were transferred into WT mice followed by PyMG tumor injection or acute LCMV- Armstrong infection. On day 8, SMARTA cells from tumor dLN or spleen of LCMV-infected mice were isolated. SMARTA cells from uninfected mice were used as naïve control. (A) Heatmap depicts the normalized z-scores of the arcsinh-transformed MSI of the indicated protein for each cluster. (B) Radar plot depicts the relative arcsinh-transformed MSI of the indicated proteins by each group, scaling as the percentage of the highest expression. (C) Heatmaps show the differential state comparisons (calculated by diffcyt (*69*)) between tumor-specific CD4 SMARTA T cells and LCMV-specific CD4 Th1 (top) or Tfh (bottom) SMARTA T cells. Coloration shows log2 fold-change. *, adjusted p-value < 0.05 (D) Violin plots show the single cell scoring of each metabolic pathway calculated as described in method. Statistics were performed Wilcox test only comparing between CD4 T_TS_ cells to Th1 and Tfh. *, p < 0.05 Data are representative of 2 experiments with at least 3 mice per group.

In contrast to glycolysis, many proteins involved in oxidative phosphorylation (OXPHOS) are increased in CD4 T_TS_ cells as compared to virus-specific CD4 Th1 and Tfh cells. For example, key enzymes involved in the tricarboxylic acid (TCA) cycle, such as citrate synthase (CS) and isocitrate dehydrogenase 1(IDH1), and mitochondrial membrane ATP synthase Complex V (ATP5A) are upregulated in CD4 T_TS_ cells (Figure 3A-C, S4A), suggesting an enhanced reliance on OXPHOS for energy production. Interestingly, the expression of cytochrome c (CytC), a critical protein in the electron transport chain for the generation of ATP, and other mitochondria proteins (such as DRP1, OPA1 and VDAC1) that are critical for mitochondria fission and function, were reduced in CD4 T_TS_ cells (Figure 3A-C, S4A), suggesting abnormal mitochondria function. Thus, CD4 T_TS_ cells show decreased mitochondria function as compared to virus-specific Th1 and Tfh cells, which could ultimately offset the enhanced TCA cycle and further result in deficiency in energy production.

Consistent with increased expression of proteins involved in the TCA cycle, CD4 T_TS_ cells have elevated amounts of enzymes involved in glutamine metabolism, whose products directly fuel OXPHOS. Several key enzymes involved in glutamine metabolism, such as glutaminase (GLS), GOT2 and glutamate dehydrogenase 1 and 2 (GLUD1/2) were significantly elevated in CD4 T_TS_ cells (Figure 3A-C, S4A). Interestingly, elevated level of GLS and glutamine metabolism has been reported to inhibit Th1 differentiation(*30*), suggesting a possible role of enhanced glutamine metabolism in the inhibition of Th1 differentiation of CD4 T_TS_ cells. The enzymes involved in lipid metabolism exhibited more complex regulation patterns. For example, medium-chain acyl-CoA dehydrogenase (ACADM) had increased levels in CD4 T_TS_ cells, whereas CPT1A was decreased (Figure 3A-C, S4A). Intriguingly, PPARγ, an important protein regulating the uptake of fatty acids and cholesterol balance in adipocytes and macrophages (*31–34*) was minimally upregulated in CD4 T_TS_ cells compared to naïve cells and was diminished compared to virus-specific CD4 Th1 and Tfh cells (Figure 3A-C, S4A). NRF1, an endoplasmic reticulum protein that directly senses cholesterol accumulation (*35*), was elevated in CD4 T_TS_ cells (Figure 3A-C, S4A). Altogether, the abnormal expression of PPARγ and NRF1 suggest a dysregulation of lipid metabolism, especially cholesterol, in CD4 T_TS_ cells.

To evaluate the overall changes of each metabolic pathway, we calculated the scores of each metabolic pathway on the single cell level by summarizing the expression of key enzymes directly involved in each pathway, as described in (*25*). Indeed, CD4 T_TS_ cells showed lower scoring in glycolysis, and significantly higher scoring in glutamine and OXPHOS (Figure 3D). Thus, CD4 T_TS_ cells exhibit signs of failure to transition to aerobic glycolysis, and instead appear to have a preference to skew toward OXPHOS despite an abnormal mitochondria metabolic state that could ultimately impair the ATP production.

### CD4 T_TS_ cell paralysis is associated with a distinct transcriptional programming in mouse and human

To understand the programming guiding CD4 T_TS_ cell differentiation, we performed single-cell (sc) RNA-seq on SMARTA CD4 T_TS_ cells from the dLN on day 8 after PyMG initiation. Clustering the CD4 T_TS_ cells revealed seven clusters (Figure 4A). Cluster 6 (enriched with activation-associated genes *Egr1,2,3, Rel, Nr4a1, Nfatc1, Cd40l and Id3*) and c7 (enriched with interferon-stimulated genes *Ifit1, Ifit3, Isg15, Isg20, Bst2, Stat1 etc*) each constituted less than 5% of the sample and were not further characterized in the following analysis (Figure 4B). Cluster 3 expressed the highest levels of *Sell* (encoding CD62L) and *Il7r*, indicating these as the naïve-like population of CD4 T_TS_ cells (Figure 4B). However, c3 also exhibited increased expression of *Jun* and *Fos* (together forming the AP-1 signaling complex)(*36*) and *Cd69*, a gene stimulated early following TCR engagement (Figure 4B), indicating that these cells have encountered antigen, but were frozen in a naïve-like state. Cluster 4 and c5 were enriched in genes involved in cell cycling. c4 was specifically enriched in protein translation (*Eif5a, Ddx21* etc), ribosome biosynthesis (*Ncl, Nop58, Pa2g4, Fbl etc*), and mitochondria function (*C1qbp, Hspd1, Atp5g1etc*), while c5 was enriched for genes involved in proliferation (*Mki67*, *Birc5, Ccnb2, Cenpe etc*) and DNA replication (*Pclaf, Top2a etc*) (Figure 4B), indicating that c4 and c5 are in different phases of cell cycle. The largest activated populations, c1 and c2 (comprising over 50% of the CD4 SMARTA T_TS_ cells) were enriched in genes that diminish T cell activation, including *Izumo1r* (FR4, associated with T cell anergy) (*37*), *Rgs10* (restricts T cell adhesion and integrin expression) (*38*) and *Pdcd4* (restricts T cell in vitro expansion) (*39*) (Figure 4B). Interestingly, recent studies have suggested that *Pdcd4* transcription is induced by CTLA4 (the most upregulated inhibitory receptor on the CD4 T_TS_ cells; Figure 2F) via FOXO1 nuclear translocation(*40*), suggesting an involvement of CTLA4 in the paralysis of these CD4 T cells. Cluster 2 was also enriched in negative regulatory genes, such as *Lgals1* (Galectin1) (*41, 42*), *Pycard/Asc* (*43*) and *Klf2* (associated with naïve T cells and quiescent state) (*44, 45*) (Figure 4B), suggesting that these cells have passed through the activation state of c1 and were suppressed into a resting state.

**Figure 4.**
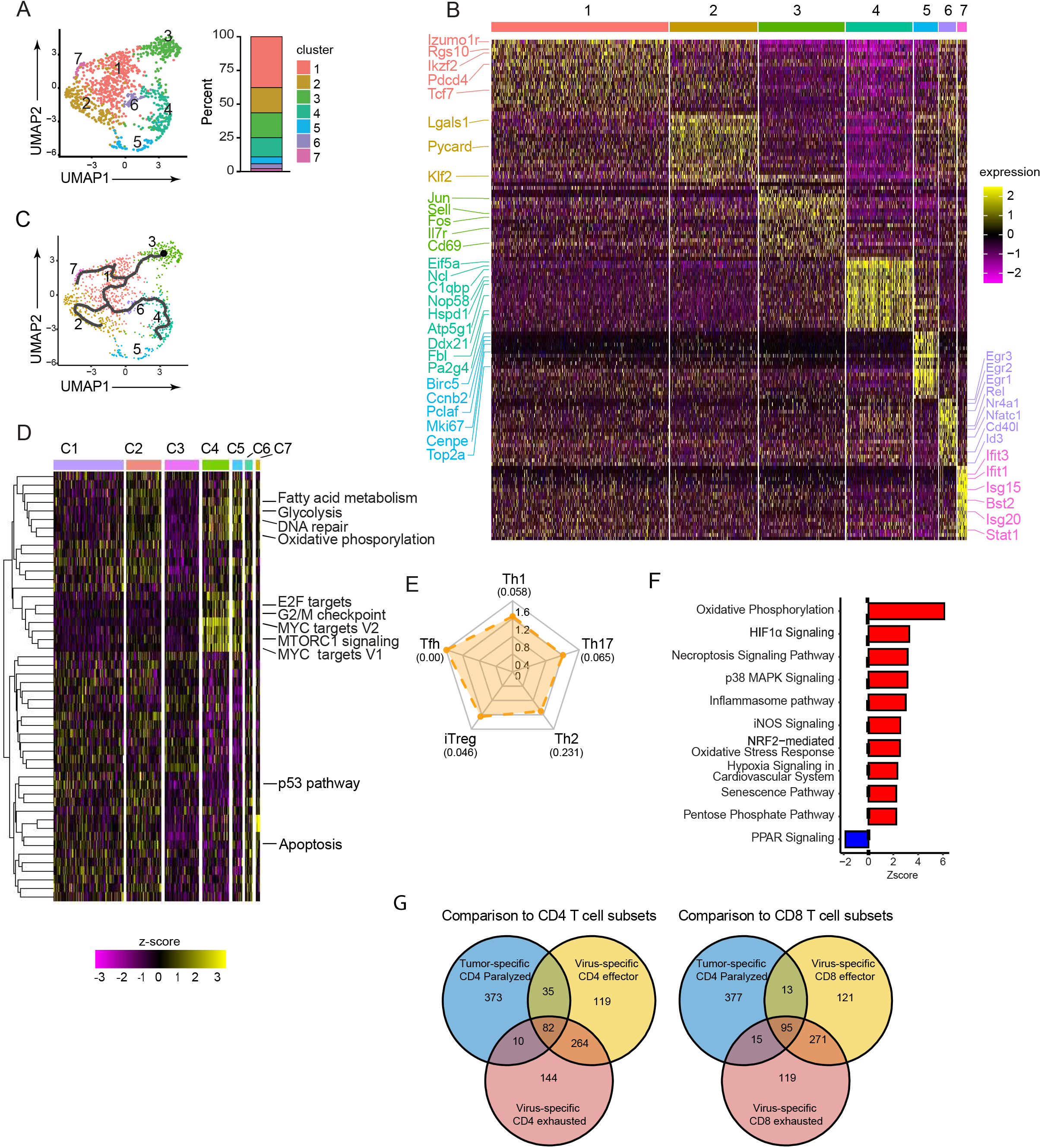
Transcriptional programming of CD4 T cell paralysis in dLN by scRNA-seq. (A) Seurat clustering (resolution 0.5) of CD4 T_TS_ cells. The bar graph depicts the proportion of each cluster. (B) Top 20 differentially-expressed genes (DEGs) of each cluster. (C) Monocle trajectory analysis overlaid on UMAP colored by Seurat clusters. (D) Single cell heatmap depicts GSVA pathway analysis within each Seurat cluster. (E) Radar plot shows normalized enrichment scores (NES) of GSEA comparing CD4 SMARTA T_TS_ cells from c4 (See Figure S4E) to defined Th cell lineage signatures. Numbers in parenthesis are the FDR. A value of 0.00 = FDR<10^-3^. (F) IPA pathway enrichment scoring of CD4 SMARTA T_TS_ cells in c4 (See Figure S4E). All included pathways are significant with p<0.05. Pathways colored in red were specifically mentioned in the text. (G) Venn diagrams show the overlap and difference in DEGs of paralyzed tumor CD4 T_TS_ cells, effector and exhausted virus-specific CD4 (left) and CD8 T cells (right). Each group was individually compared to naïve T cells first prior to the comparisons shown. The numbers indicate the number of genes.

We performed Monocle (*46*) analysis to determine the differentiation trajectory of the CD4 T_TS_ cells. Starting from the naïve c3, the differentiation trajectory goes through a common population (c1) then forks into 3 distinct paths (Figure 4C). Consistent with c1 as the first path through which the other populations progress, cells in c1 had the lowest differentiation score, a score that ranks pseudotime distance from the starting population (*46–48*) (Figure S4B). The first fork from c1 leads to a pathway enriched in interferon stimulated cells (c7). At the second fork, one direction path leads to c2, with these cells having an intermediate pseudotime score (Figure S4B), consistent with their increased expression of negative regulatory genes compared to c1 (Figure 4B). The third path leads to actively dividing populations (c4 and c5) with the highest pseudotime scores (Figure S4B), suggesting that a higher differentiation state is associated with active cell cycle. Thus, most CD4 T_TS_ cells are stuck in an early-priming stage by day 8, upregulating genes that antagonize activation; and although a small population are proliferating, the majority express activation-suppressing genes.

We next preformed gene set variation analysis (GSVA) to investigate the transcriptional pathways associated with each population. Consistent with their proliferation status, c4 and c5 highly upregulated multiple gene expression pathways involved in cell cycling, such as E2F targets, G2/M checkpoints and DNA repair, whereas naive-like population (c3) and the paralyzed populations (c1 and c2) did not (Figure 4D, and all differentially expressed pathways in each cluster are quantified in S4C). As expected, the actively proliferating cells (c4 and c5) had undergone extensive metabolic reprogramming, particularly upregulating glycolysis in parallel to high activation of the Myc pathway (required for effector T cell metabolic reprogramming). Consistent with its low differentiation status, c1 is metabolically lower for gene pathways supportive of glycolysis, OXPHOS and fatty acid metabolism, and this population fails to activate Myc pathway (Figure 4D, S4C). Although c2 upregulated fatty acid metabolism and OXPHOS (albeit lower than the actively proliferating populations c4 and c5), c2 did not significantly upregulate glycolysis or activate Myc (Figure S4C). Interestingly, compared to proliferating cells (c4 and c5), both c1 and c2 exhibited low expression of the mTORC1 signaling pathway, a major cell-activation pathway induced by TCR signaling (*49*) (Figure 4D, S4C), suggesting that the proliferative paralysis and diminished metabolic reprogramming could be a result of insufficient priming. In contrast to the metabolic and proliferation pathways, apoptosis and P53 pathways (involved in cell cycle arrest) are activated in c1 and further enhanced in c2, but not in naïve-like population (c3) or proliferating cells (c4 and c5) (Figure 4D, S4C). Altogether, these observations suggest that priming by tumor antigen fails to induce necessary cellular changes essential for sustained clonal expansion of effector CD4 T cells.

To define the global changes of T cell paralysis, we re-clustered with naïve CD4 T cells. This approach yielded a dominant cluster (c4) in the cells from the tumor-bearing mice and multiple clusters in the naïve cells. The dominant clusters in the CD4 T_TS_ cells and naïve cells were generally exclusive of each other (Figure S4D). For the analysis, the naïve cells were combined, only excluding a small activated cluster 7 (Figure S4D). Compared to naïve CD4 T cells, CD4 T_TS_ cells exhibited a mixed Th transcriptional programming, featuring enhanced Tfh and to a lesser extent iTreg signatures, with limited Th1, Th2 and Th17 signatures (Figure 4E), consistent with the protein expression analysis by CyTOF (Figure2A, B). Ingenuity pathway analysis (IPA) further revealed that OXPHOS and pentose phosphate pathways were activated in CD4 T_TS_ cells but not glycolysis (Figure 4F). Consistent with the low protein expression (Figure 3A-C, S4A), upstream regulator analysis indicated inhibited activity of HK2 (glycolysis) and PPAR pathways (Figure 4F, S4E), further corroborating a defect in glycolysis and PPARγ-regulated fatty acid metabolism and intracellular cholesterol balance. In addition, CD4 T_TS_ cells are enriched in pathways associated with cellular stress, including NRF2-mediated oxidative stress, hypoxia and HIF1α signaling (Figure 4F). In parallel to the oxidative stress response, pathways negatively related to survival and proliferation, such as necroptosis, senescence and inflammasome signaling, are also activated in these CD4 T_TS_ cells (Figure 4F), whereas the activity of transcription factor ID2 (important for effector T cell survival and resistance to apoptosis) is inhibited (Figure S4E). Thus, CD4 T_TS_ cells fail to undergo necessary metabolic reprogramming, and exhibit abnormal cellular stress signaling and pathways negatively associated with survival.

Next, we compared the gene expression profiles of CD4 T_TS_ cells to previously defined effector and exhaustion gene expression signatures (*50–52*). As a first step, all populations were individually compared to naïve T cells to generate up-regulated genes specific to each cell state. Comparing those up-regulated genes against gene expression profiles of effector and exhausted CD4 T cells demonstrated that most up- regulated genes in the CD4 T_TS_ cells are unique to that subset, with 75% of the top 500 up-regulated genes in the CD4 T_TS_ cells being unique compared to effector or exhausted CD4 T cells (Figure 4G). Further, the paralyzed CD4 cells have minimal overlap of with CD4 effector (only 12% of DEGs overlap) or exhausted (9% of DEGs overlap) T cells. The same lack of overlap is observed in relation to CD8 effector (11% overlap of up-regulated genes) or exhausted (11% overlap of up-regulated genes) T cells (Figure 4G). Thus, combined with the CyTOF and transcriptional data, the CD4 T cell paralysis is a cellular state distinct from T cell effector or exhaustion.

To investigate whether a similar population of CD4 T cells was present in humans, we interrogated published scRNA-seq data from the tumor dLN of patients with breast cancer (*53*) and lung adenocarcinoma (*54*). For this analysis, we used the 373 DEGs identified in Figure 4G as the CD4 T_TS_ signature. Up to 8.3% of sequenced human dLN CD4 T cells possessed a similar gene expression to mouse paralyzed CD4 T_TS_ cells (Figure S4F, Table S4). Interestingly, the number of this population increased with lymph node metastasis. We then compared these human CD4 T cells against the remaining CD4 T cells in the sample to identify a specific signature associated with human paralyzed CD4 T cells (Figure S4G). Biological processes GO enrichment analysis of these signature genes demonstrated that the human CD4 T cells were enriched in similar pathways as paralyzed mouse CD4 T_TS_ cells, particularly those related to oxidative phosphorylation (Figure S4H). Thus, a small population of CD4 T cells in dLN of human breast and lung cancer patients also exhibit a similar CD4 T_TS_ cell paralysis transcriptional signature.

### CTLA4 blockade alleviates the proliferative paralysis but fails to promote the differentiation of effector CD4 Th subsets

We next investigated the molecular mechanisms underlying the CD4 T_TS_ cell proliferation and differentiation paralysis. Both CTLA4 and to a much lesser extent PD1 were upregulated by CD4 T_TS_ cells early after activation (Figure 2F), suggesting a role for these inhibitory receptors. Consistent with its minimal expression, blocking PD1:PDL1 interactions during initial activation failed to overcome the proliferative paralysis (Figure S5A). In contrast, CTLA4 blockade completely restored the proliferation of CD4 T_TS_ cells (Figure 5A). Further, blocking CTLA4 in mice with established tumors (day 21) also restored CD4 T_TS_ cell proliferation (Figure 5B), indicating that CTLA4 constantly maintains proliferative paralysis through tumor progression. To further investigate how CTLA4 blockade alone impacts CD4 T_TS_ cell response, we used CyTOF to characterize SMARTA cells in day 8 dLN of CTLA4 blockade or isotype mice. Despite promoting proliferation, CTLA4 blockade only induced the emergence of a single cluster (c8), with minimal effect on protein expression, except for increased ICOS (Figure S5B-D), suggesting an uncoupling of proliferation from differentiation in CD4 T_TS_ cells. The newly emergent c8 following CTLA4 blockade did express increased activation-induced proteins and appeared to be a mixed Th1-Tfh precursor (based on Tbet and Bcl6 co-expression) (Figure S5C). Both Tregs and tumor-specific CD4 T cells express CTLA4 in the dLN. Inducible deletion of CTLA4 expression on Tregs (FoxP3-icre x CTLA4fl/fl mice) prior to tumor implantation did not overcome the proliferative paralysis (Figure S5E), nor did CRISPR-mediated deletion of CTLA4 in SMARTA cells prior to transfer (Figure S5F), indicating a cooperative mechanism of CTLA4 on both Treg and CD4 T_TS_ cells to inhibit proliferation.

**Figure 5.**
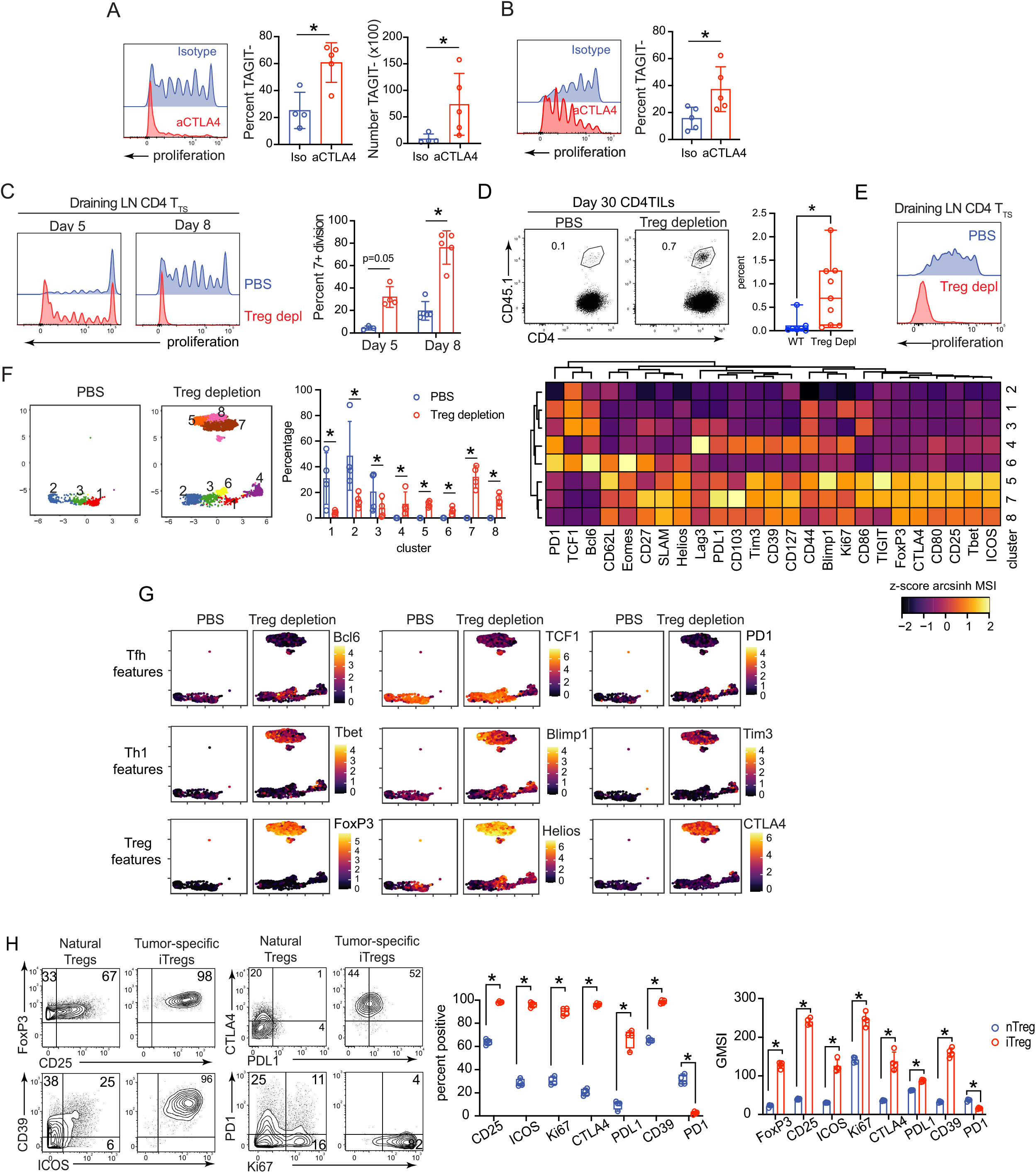
CTLA4 and Tregs reciprocally induce paralysis of CD4 T_TS_ cells. (A, B) On days 0, 2 and 5 (A); or days 21, 23 and 26 (B) after PyMG administration, mice were treated with 300ug CTLA4 blocking or isotype control antibodies. Histogram shows proliferation of SMARTA cells on (A) day 8 or (B) day 29 in the dLN. Bar graphs show the frequency and number of SMARTA cells that completely diluted TAGIT. (C-H) FoxP3-DTR mice received TAGIT labeled naïve SMARTA cells followed by PyMG injection and either PBS or diphtheria toxin (DT) treatment. (C) Histograms show the proliferation of CD4 SMARTA T_TS_ cells in the dLN on the indicated day. The bar graph shows the percentage of CD4 SMARTA T_TS_ cells that have completely diluted TAGIT. (D) Flow plots and bar graph depict the frequency of SMARTA cells of CD4 T cells in the tumor at day 30. (E) FoxP3-DTR mice were treated with PBS or DT on day 21 and 23 after PyMG injection. Histograms show the proliferation of CD4 SMARTA T_TS_ cells in dLN on day 29 (8 days after treatment). (F) UMAP plots indicating PhenoGraph clusters of CD4 SMARTA T_TS_ cells in dLN 8 days after PyMG initiation (concatenated from 4-5 mice in each group). Bar graph shows the frequency of each cluster, with each circle representing the percent of SMARTA T_TS_ cells in that cluster from an individual mouse. * = adjusted p-value <0.05, calculated by diffcyt. Heatmap depicts the normalized z-scores of the arcsinh- transformed MSI of the indicated protein for each cluster (G) UMAPs show the single cell arcsinh-transformed expression of the indicated protein by CD4 SMARTA T_TS_ cells. (H) Expression of indicated protein on SMARTA iTreg (c5, 7 and 8) and Tregs (FoxP3+ Helios+ of non- SMARTA CD4 T cells in dLNs, from PBS control treated mice). Bar graphs show the frequency (left) and GMSI (Geometric mean of arcsinh-transformed MSI calculated by FlowJo^TM^) (right) of the indicated protein in the Tregs or SMARTA iTregs. Data represents 2-3 independent experiments with at least 4 mice per group. Error bars indicate SD. Significance other than CyTOF analyses is determined by Mann-Whitney U-test. *, p<0.05.

### Tregs mediate the CD4 T cell proliferative paralysis, but their depletion reciprocally induces tumor-specific iTreg differentiation

Based on the expression of CTLA4 by Tregs, we next sought to understand how Tregs were modulating the CD4 T cell paralysis. Coinciding with the CD4 T_TS_ cell paralysis was an increase in the frequency of FoxP3+ Tregs in the dLN (Figure S6A). Indeed, depleting Tregs at the time of tumor initiation using mice expressing diphtheria toxin receptor (DTR) under the control of the FoxP3 promoter (FoxP3- DTR; DREG) led to rapid CD4 T_TS_ cell proliferation in the dLN (Figure 5C, S6B) and sustained tumor infiltration (Figure 5D). A similar increased proliferation occurred when Tregs were depleted using anti- CD25 antibody (Figure S6C). Further, Treg cell depletion in mice with established tumors (21 days after PyMG initiation) overcame the proliferative paralysis and induced robust CD4 T_TS_ cells division (Figure 5E, S6D). Thus, Treg cells actively and continuously inhibit CD4 T_TS_ cell proliferation throughout tumor progression.

In addition to overcoming the proliferative block, Treg depletion increased the activation state of CD4 T_TS_ cells and drove the emergence of multiple new CD4 T_TS_ cell clusters (c4-8) (Figure 5F). Of the newly emerging cell populations after Treg depletion, c6 exhibited protein-expression patterns of Tfh-like cells with high expression of Bcl6, TCF1, and PD1, while retaining high CD62L (Figure 5F, 5G). Conversely, c4 differentiated into Th1-like cells (a subset previously unseen when Tregs were present), expressing Tbet, Blimp1, Tim3 and Lag3, combined with low expression of TCF1 and Bcl6 (Figure 5F, 5G). Further, both the Th1-like c4 and the Tfh-like c6 had higher expression of CD27, CD127 and Ki67 (Figure S6E), suggesting increased activation and survival. Strikingly, Treg depletion induced the formation of multiple clusters with Treg phenotypes (c5, 7 and 8) that now comprised ∼50% of the total CD4 T_TS_ cells (Figure 5F) that was not observed after CTLA4 blockade (Figure S6F). These tumor-specific iTregs expressed high levels of FoxP3, Helios and CTLA4 (Figure 5F, 5G), with a subset expressing both CD69 and CD103 (Figure S6G), suggesting increased tissue homing/retention capacity. When compared to the Tregs in PBS-treated mice, the tumor-specific iTregs had increased expression of FoxP3 and multiple functional proteins (CD25, CD39, ICOS, CTLA4, PDL1), Th1-associated proteins (SLAMF1, Blimp1, Tbet and Tim3), but lack TCF1, Bcl6 and did not express PD1 (Figure 5G, 5H, S6H). This increased activation-protein expression was evident in the frequency of cells expressing these proteins and at the single-cell expression level in the tumor-specific iTreg (Figure 5H, S6H). Thus, Tregs rapidly induce and then maintain the CD4 T_TS_ cell paralysis, and their depletion reciprocally promotes the differentiation of highly-activated tumor-specific iTregs.

Mechanistically how CTLA4 blockade and Treg depletion impair CD4 T cell proliferation in the dLN is unclear. Untreated CD4 T_TS_ cells exhibit a transcriptional and metabolic profile consistent with sub-optimal activation. Indeed, depleting Treg or CTLA4 blockade specifically led to increased number and heightened activation of a specific population of DCs (a CD11b+ DC2 pop I) in the dLN, with elevated expression of the co-stimulatory molecules CD80 and CD86 (Figure S6I-J). Surprisingly, Treg depletion significantly increased MHCII expression on these cDC2 population whereas CTLA4 blockade did not (Figure SI-J). These observations suggest that Tregs and CTLA4 synergistically induce T cell paralysis by limiting signals for T cell activation. Indeed, treating mice with aCD3 and/or aCD28 agonists antibodies successfully restored the proliferation of CD4 T_TS_ cells in dLN in the presence of Tregs and CTLA4 (Figure 6A). Thus, the Treg and CTLA4 mediated paralysis of CD4 T_TS_ cells is overcome by increasing T cell activation signaling.

**Figure 6.**
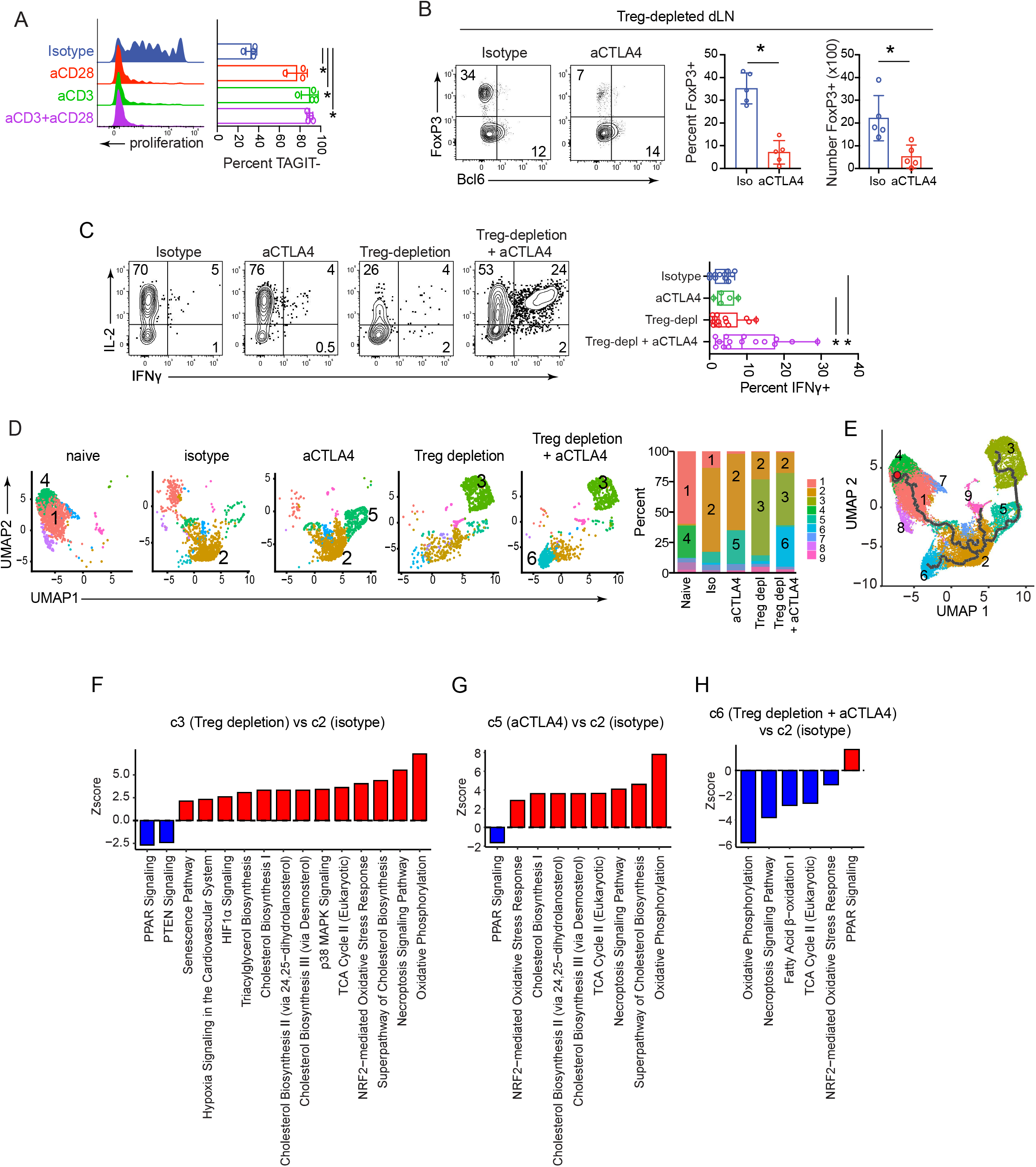
CTLA4 drives tumor-specific iTreg differentiation in Treg-depleted mice and dual Treg depletion and CTLA4 blockade enable effector differentiation and metabolic re-organization. (A) On day 5 after PyMG administration, mice received isotype control, anti-CD3 alone, anti-CD28 alone, or dual anti-CD3 and anti-CD28 antibodies. Cell proliferation of CD4 SMARTA T_TS_ cells in dLN was analyzed on day 8. (B-C) FoxP3-DTR mice were treated on days 0 and 2 with DT, and on days 0, 2 and 5 with CTLA4 blocking or isotype antibodies. (B) Expression of FoxP3 and Bcl6 by SMARTA cells. Bar graphs show the frequency (left) and number (right) of FoxP3+ SMARTA cells. (C) Cytokine production by CD4 SMARTA T_TS_ cells after *ex vivo* restimulation. (D-H) FoxP3-DTR mice were treated on days 0 and 2 with PBS or DT; and / or on days 0, 2 and 5 with CTLA4 blocking or isotype antibodies. Analysis was performed on dLN SMARTA T_TS_ cells on day 8 after PyMG administration. (D) UMAP plots show Seurat clustering (resolution 0.3) of CD4 SMARTA T_TS_ cells s in each condition. Bar graph depicts the proportion of each cluster. The numbers in the stacked bar graphs indicate the cluster. (E) Monocle Trajectory analysis overlaid onto the Seurat clustered UMAP (all conditions combined). (F-H) IPA pathways enrichment scoring of the indicated clusters. All pathways are significant with a p-value < 0.05. For Panels A-C, the data represent 3 independent experiments with at least 5 mice per group. Error bars indicate SD. Significance is determined by Mann-Whitney U test. *, p<0.05.

### Distinct effects of CTLA4 blockade and Treg depletion on tumor-specific CD4 T cell differentiation and programming

Since CTLA4 blockade did not induce SMARTA iTreg differentiation as Treg depletion, we blocked CTLA4 signaling in combination with Treg depletion. The combinatory treatment diminished the *de novo* iTreg differentiation (Figure 6B), indicating that CTLA4 signals on the CD4 T_TS_ cells induce the iTreg differentiation when the suppression from Tregs is alleviated. In addition to reduced iTreg differentiation, the combination treatment significantly enhanced the production of IFNγ, while slightly decreasing IL-2 and TNFα production (Figure 6C, S6K). To understand the changes in cellular transcriptional networks underlying CTLA4 and Treg mediated suppression of CD4 T_TS_ cells, we performed scRNA-seq. Tumor-specific CD4 SMARTA T cells were isolated from the dLN on day 8 after PyMG initiation following: (A) isotype antibody treatment, (B) CTLA4 blockade, (C) Treg depletion, or (D) dual Treg depletion with CTLA4 blockade. Naïve SMARTA cells were used for comparison. Seurat clustering revealed nine clusters across the conditions, with certain clusters emerging only following specific treatments (Figure 6D). Three minor clusters (c7-9) each constituted less than 5% in each condition and their frequencies remained nearly unchanged after each treatment, therefore they were not included in further analysis. Naive SMARTA cells were composed of two similar populations based on their cluster defining genes (c1 and c4) with high levels of *Ccr7, Il7r and Tcf7*, and therefore were combined in the future pathway analysis (Figure S7A). The majority of CD4 T_TS_ cells in the dLN from isotype treated mice formed a main c2 expressing a similar gene signature to our previous analysis (Figure 4B), such as high expression of *Izumo1r* (FR4), *Rgs10* and *TCF7* (Figure 6D, S7A). Treg depletion led to an increase in iTreg (c3), characterized by *FoxP3* and *Ikzf2* (Helios) and other Treg-associated genes, such as *Tnfrsf4* (OX40) and *Ctla4* (Figure 6D, S7A), further confirming the iTreg differentiation. Blocking CTLA4 alone induced a unique c5 that, although it did not express the Treg lineage defining genes (*Foxp3, Ikzf2, Ctla4*), it shared many DEGs with the iTreg c3, including those involved in DNA transcription and repair, including *Hmgb2*, *Hmgn2* and *Pclaf* (Figure 6D, S7A). The frequency of the iTreg c3 was reduced when blocking CTLA4 and depleting Tregs (in this case by ∼20%), and a new c6 developed, characterized by high expression of genes associated with T cell activation and function, including *Fos*, *Fosb*, *Jun* (together forming the AP-1 complex) (*36*), *Nkg7* (lymphocyte granule exocytosis) (*55*), *Lag3*(*56, 57*) and *Eomes* (*58*) (Figure S7A), consistent with the CyTOF data demonstrating combination treatment promoting differentiation of highly activated CD4 T_TS_ cells. Cell fate trajectory analyses revealed that c5 (the population that emerged only in CTLA4 blockade alone) and c3 (iTregs that emerged only in Treg depletion alone) are on the same differentiation pathway, whereas c6 (the population that emerged in dual treatment) had a distinct differentiation pathway (Figure 6E). Interestingly, the newly emergent clusters in the various treatment modalities all progress through c2 (Figure 6E), consistent with this being the initial path of differentiation that CD4 T_TS_ cells are subsequently frozen in by CTLA4 and Tregs.

We next used IPA to perform pathway and upstream regulator analyses comparing c2 (the main cluster in isotype treated mice) with the emergent clusters in each condition: c3 (the population that emerged only in Treg-depleted), c5 (the population that emerged only in anti-CTLA4), and c6 (the population that emerged only in dual treatment). Among the top upstream regulators activated by either treatment alone are those associated with TCR and costimulation, including CD3, CD28, LCK, MTORC1, NFkB complex and PI3K complex (Figure S7B), thus further corroborating that both Treg and CTLA4 diminishes antigen signaling to paralyze the CD4 T_TS_ cells. The newly emergent cluster c3 after Treg depletion alone or c5 following CTLA4 blockade alone further elevated the OXPHOS metabolism pathway, particularly the TCA cycle, compared to isotype (Figure 6F, 6G); while IPA identified the glycolytic enzymes GAPDH and HK2 as upstream regulators that were even further inhibited (Figure S7B). Lipid biosynthesis, particularly cholesterol and triacylglycerol biosynthesis, is further promoted by CTLA4 blockade or Treg depletion alone, along with the further inhibition of the PPAR pathway (Figure 6F, 6G). In contrast, dual CTLA4 blockade and Treg depletion inhibited OXPHOS and cholesterol biosynthesis pathways and restored PPAR activity compared to the other conditions (Figure 6H, S7C). Thus, even though single treatments alone promoted proliferation (by enhancing T cell stimulation and likely through increased energy production), it exacerbated the reliance on OXPHOS and cholesterol biosynthesis, which are only corrected by the combination treatment. In addition to metabolic changes, the combination treatment also induced gene expression pathways that promote survival. Either CTLA4 blockade or Treg depletion alone exacerbated the activation of oxidative stress, as exemplified by the activation of upstream regulator pathways NRF2 and HIF1a, and pathways that negatively affect cell survival, such as necroptosis and senescence (Figure 6F, 6G, S7B) that were already activated in untreated tumor-activated CD4 T cells (Figure 4F). In contrast, these pathways are inhibited in cells receiving dual treatment as compared to isotype sample (Figure 6H) or compared to single treatment sample (Figure S7C). Thus, dual CTLA4 blockade and Treg depletion have combined effects to improve cellular metabolism, reduce oxidative stress, and promote survival and further differentiation.

## Discussion

We demonstrate that dLN-resident CD4 T_TS_ cells are primed and begin to divide following tumor initiation. However, unlike CD8 T_TS_ cells that continue to proliferate and accumulate in the tumor, the proliferation of CD4 T_TS_ cells is rapidly frozen in place, stunting their differentiation and limiting their accumulation in the tumor to only a small subset of CD4 TTS cells that had extensively divided. This is consistent with the observation from the clinical studies that the tumor is extensively infiltrated by “by-stander” T cells (*12–14, 59*).The paralysis is accompanied by gradual cell death over-time, which would further limit the quality of the CD4 T_TS_ cell response and suggest that early interventions that could amplify the response prior to attrition would be most beneficial. As a result, the help provided by CD4 T_TS_ cells is rapidly limited and suboptimal for tumor control. Over time, a fraction of the paralyzed CD4 T cells are converted to iTregs in both dLN and the tumor, suggesting that the tumors take advantage of the small amount of residual tumor-specific CD4 T cell differentiation to further promote their growth. The proliferative paralysis and stunted Th differentiation were actively maintained throughout tumor progression. Yet, CD4 T_TS_ cells retained the ability to resume proliferation and differentiation when the suppressive constraints were abrogated either early or late in tumor progression, demonstrating that the tumor actively and continually paralyzes CD4 T cells as a mechanism to promote cancer cell growth. Overcoming the proliferative paralysis with bmDCs administration led to enhanced tumor control at the beginning and in the established tumor, highlighting the importance for the tumor to rapidly inhibit and then maintain CD4 T_TS_ cell paralysis.

Rather than experiencing functional exhaustion in the TME that is driven by heightened / prolonged antigenic stimulation and PD1/PDL1 interaction, the CD4 T_TS_ cell response is restricted early following initial activation through multiple mechanisms that rapidly quell activation (Figure S8), although the CD4 TTS that infiltrate the tumor may then be restricted by T cell exhaustion. During priming, cooperative pathways are engaged to limit TCR and co-stimulatory molecule signaling at the level of the CD4 T_TS_ cells and the APC, resulting in the paralyzed state. In the presence of Treg cells, DCs express lower levels of T cell activating components, including MHCII and B7 family molecules. This is accompanied by the upregulation of CTLA4 on the CD4 T_TS_ cells that further limits TCR signal and competes for binding to CD80 and CD86. Together, these signals collaborate to diminish T cell activation. Alleviating either of these inhibitors of T cell activation (or exogenously overcoming them with antibody-mediated antigenic stimulation) restored TCR signaling and costimulation to resume proliferation. Counterintuitively, depleting Treg alone reciprocally induced highly-activated tumor-specific iTreg to take their place. Interestingly, CTLA4 played a dual role in this process. Initially, CTLA4 inhibited TCR/costimulation signals to prevent proliferation. Then following Treg depletion, CTLA4 provided the signals to drive iTreg differentiation. Only by simultaneously alleviating both Tregs and CTLA4 could the differentiation of effector Th cells and the production of IFNγ be induced. It was interesting that the restricted CD4 Th differentiation included limited PD1 upregulation. PD1 blockade has been shown to minimally affect CD4 T cells in tumor models (*7*), and only specific Th subsets are amenable to PD1 mediated functional, specifically Th1 and Treg, with limited effect on Tfh (*9*). Thus, the low expression of PD1 and uncommitted Th state may explain why PD1 blockade fails to restore CD4 T_TS_ cells and preferentially enhances Treg responses in these cancer models.

The regulation of CD4 T_TS_ cells occurred early within the dLN. CD4 T_TS_ cells that did enter the tumor had extensively proliferated, suggesting that full proliferation is required for egress from the dLN to the tumor. CTLA4 blockade and Treg depletion restored the proliferation of CD4 T_TS_ cells in dLN, and again in this situation, the CD4 T_TS_ cells that infiltrated the tumor had completely diluted proliferation dye. This is consistent with clinical observations and mouse models that immune-checkpoint blockades induce T cell clones previously not observed within the tumor (*60*). Our study provides one explanation for this observation by demonstrating that CD4 T_TS_ cells are diminished in the dLN and especially the tumor due to their restricted proliferation. By expanding CD4 T_TS_ cells from their low levels, immunotherapy enables their homing to the tumor where they can become identifiable clones. Thus, in addition to inducing de novo tumor-reactive CD4 T cell priming, immunotherapy can also expand low amounts of pre-existing paralyzed clones to home to the tumor. In essence, it is not that these clones did not exist before, but only became quantifiable following their therapy-induced expansion. Further, our study highlights the importance of spatial restriction of CD4 TTS cells to the dLN as a reservoir for restorative potential. Thus, we demonstrate a new mechanism of CD4 T cell proliferative, functional and metabolic suppression that promotes tumor immune evasion.

## METHODS

### Mice

For all experiments, female mice were used for breast cancer cell lines (PyMT and PyMG) and either female or male mice were used for B16-F10 and colorectal carcinoma (MC38 and MC38GP). C57BL/6 mice (7-12 weeks old) were purchased from Princess Margaret Cancer Center or The Jackson Laboratory. LCMV-GP_61-80_-specific CD4 TCR transgenic (SMARTA; CD45.1+) mice and LCMV-GP_33-41_-specific CD8 TCR transgenic (P14; CD45.1+) were described previously (*61*). OT-II mice (Stock No: 004194) were purchased from The Jackson Laboratory. CD4KO (Stock No: 002663) and FoxP3-DTR (Stock No: 016958) breeders were purchased from The Jackson Laboratory and maintained in the mice facility at Princess Margaret Cancer Center. β_2_m-/- female mice (Stock No:002087) were purchased from The Jackson Laboratory and used for harvesting bone marrow. CTLA4 flox/flox mice were provided by Dr. Pamela Ohashi (University health Network) and crossed to FoxP3^eGFP-Cre-ERT2^ (Jackson stock No: 016961). All mice were housed under specific pathogen-free conditions. Mouse handling conformed to the experimental protocols approved by the OCI Animal Care Committee at the Princess Margaret Cancer / University Health Network.

### T cell adoptive transfer, TAGIT labelling and cell proliferation quantification

CD4 SMARTA T cells, CD4 OT-II T cells or CD8 P14 T cells were isolated from the spleens of transgenic mice using EasySep^TM^ mouse naïve CD4 T cell negative isolation kit (Catalog No. 19852, STEMCELL) or EasySep^TM^ mouse naïve CD8 T cell negative isolation kit (Catalog No. 19858, STEMCELL), respectively. For tumor experiments, 100,000 SMARTA, OT-II or P14 T cells were transferred to mice i.v. via retro-orbital sinus one to two days prior tumor injection. For LCMV experiments, 3,000 SMARTA cells were transferred to mice i.v. via retro-orbital sinus one to two days prior infection.

Where indicated, transgenic T cells were labeled with Tag-it Violet Proliferation and Cell Tracking Dye (Cat No. 425101 Biolegend). The frequency of SMARTA cells that proliferated during culture was determined as described by Gett et al. (*62*) . Briefly, the number of cells in each division peak was divided by 2^i^ (where i equals the number of divisions), and the resulting number of each division was then summed as the number of SMARTA cells that had divided. This number was then divided by the sum of the divided SMARTA and the number of undivided cells. This number was then expressed as the percentage of proliferation.

### LCMV infection

For experiments where LCMV was used as control, mice were infected with 2 million plaque-forming units (PFU) of LCMV-Cl13 i.v. via the retro-orbital sinus, or two hundred thousand PFU. of LCMV-Armstrong intraperitoneally (i.p.) into mice. Virus stocks were prepared and viral titers were quantified as described previously (*61*). LCMV-specific CD4+ SMARTA T cells were isolated from the spleens of transgenic mice as described above. Then 3,000 CD45.1+ SMARTA cells (donors) were transferred i.v. into the retro-orbital sinus of naïve CD45.2+ C57BL/6 mice (recipients) that were then infected with LCMV-Cl13 or -Armstrong one day later. Spleen cells were used for analysis of virus-specific CD4 T SMARTA cells.

### Tumor cell lines and LCMV-GP1-100 vector construction

PyMT and MC38OVA cell line were provided by Dr. Pam Ohashi (University Health Network). The MC38 tumor line was provided by Dr. Daniel De Carvalho (University Health Network). B16-F10 and MC38OVA was provided by Dr. Tracy McGaha (University Health Network).

To generate PyMG cell line, LCMV-GP_1-100_ was cloned into MSCV-IRES-Thy1.1 DEST vector (*63*) (MSCV-IRES-Thy1.1 DEST was provided to Addgene by Dr. Anjana Rao, Addgene plasmid# 17442 ; http://n2t.net/addgene:17442 ; RRID:Addgene_17442) using TaKaRa In-Fusion HD Cloning Plus kits (Cat# 638917) to get MSCV-LCMVGP-IRES-Thy1.1 vector. The following primers were used to amplify LCMV-GP_1-100_ and perform infusion cloning, forward primer: CGCCGGAATTAGATCATGGGTCAGATTGTGACAAT; reverse primer: ACCGGATCCAGTCGATCAGTAATGGTGGGAGTTGTT. Then MSCV-LCMVGP-IRES-Thy1.1 vector and pCL-ECO vector (*64*) (pCL-Eco was provided to Addgene by Dr. Inder Verma, Addgene plasmid# 12371 ; http://n2t.net/addgene:12371 ; RRID:Addgene_12371) were transfected in 293 T cells with lipofectamine 3000 (Cat# L3000001, Thermo Fisher) to generate virus for transfection. The parental PyMT cell line was then transfected with the pseudotyped LCMV-GP1-100 containing virus (MOI=1) in the presence of polybrene. After viral transfection, single Thy1.1+ tumor cells were isolated by FACSorting and single cell clones generated. PyMG was selected among a tested single cell clones after injecting and tracking their growth in female C57BL/6 mice.

Transposon transduction was used to generate the MC38GP cell line. LCMV-GP_1-100_-Thy1.1 sequence was amplified from MSCV-LCMVGP-IRES-Thy1.1 vector using the following primers: forward: GCCTCTGAGGCCACCATGGGTCAGATTGTGACAATGTTTG;reverse: ATTGATCCCCAAGCTTCACAGAGAAATGAAGTCCAGGGC, and then cloned into pSBbi-Pur vector (*65*) (pSBbi-Pur was a gift from Eric Kowarz, Addgene plasmid # 60523 ; http://n2t.net/addgene:60523 ; RRID:Addgene_60523) using TaKaRa In-Fusion HD Cloning Plus kits (Cat# 638917) to get pSBbi-Pur- LCMVGP-Thy1.1 vector. Then pSBbi-Pur-LCMVGP-Thy1.1 vector and pCMV(CAT)T7-SB100(*66*) (pCMV(CAT)T7-SB100 was a gift from Zsuzsanna Izsvak, Addgene plasmid # 34879 ; http://n2t.net/addgene:34879 ; RRID:Addgene_34879) were transfected into MC38 cell line with lipofectamine 3000 (Cat# L3000001, Thermo Fisher). Thy1.1+ tumor cells were then FACs sorted into single cell clones and grown with puromycin (8ug/ml, Gibco Ref A1113803) selection media. MC38GP was selected based on the growth rate in male and female C57BL/6 mice.

### Tumor cell injection

For injection of PyMT and PyMG, tumor cells were washed thoroughly with PBS after trypsinization and resuspended at 1×10^6^ cells per 20µl in PBS. Female mice received 20µl injection at the left lower 5^th^ mammary fat pad to avoid injection into nearby lymph node. For injection of MC38, MC38GP, MC38OVA and B16-F10, tumor cells were resuspended at 2×10^5^ per 50ul PBS. Female or male mice received 50ul injection at the inner side of the leg. Inguinal lymph nodes were tumor draining lymph nodes for all tumors used.

### Tissue processing

Tumors were harvested and manually dissociated into small pieces in gentleMACS^TM^ C tubes (Miltenyi Biotec) with 5ml RPMI 1640 medium with 100U/ml Collagenase I (Thermofisher 17100017) and 10ug/ml DNase I (Sigma DN25-1G) and 2% FBS. For PyMG/PyMT tumors, protocol 37_m_TDK_2 of the gentleMACS Dissociator (Miltenyi Biotec, 130-095-937) was used. For B16 and MC38/MC38GP, protocol 37_m_TDK_1 was used. Single cell suspensions were obtained by filtering through pre-separation filters (70μm) (Miltenyi, order#130-095-823). Due to the limitation of the number of live cells recovered from day 8 tumor, all flow plots concerning day 8 TME was collected by combing individual tumor samples of each group into one sample during digestion. Inguinal lymph nodes were harvested and manually dissociated into small pieces in 1.5ml centrifuge tubes with 400ul buffer I (RPMI with 10% FBS and 1% HEPES), followed by digestion with 1mg/ml Collagenase IV from Clostridium histolyticum (C5138 Sigma) and 0.15mg/ml DNase I (Sigma) at 37 degree for 30 minutes. Lymph node single cell suspension was obtained by filtering through the strainer snap cap of Falcon^TM^ 352235 round-bottom polystyrene test tubes (Fisher scientific Cat#08-771-23). Spleens were harvested from LCMV infected mice and manually dissociated into single cell suspension with a tissue smasher.

### *In vitro* generation and peptide labeling of bone marrow derived dendritic cells

bmDCs were generated as previously described (*67*) . Briefly, 2×10^6^ bone marrow cells were cultured at 37°C in 10ml bmDC culture medium (RPMI 1640 medium with 10% FBS, penicillin/streptomycin and 2-Mercaptoethanol) with 40ng/mL GM-CSF (BioLegend). At day 3, 10 ml fresh bmDC culture medium with 40ng/ml GM-CSF was added. At day 6 and 8, 10ml media was removed and replaced with fresh bmDC culture medium with 20ng/ml (day 6) or 10ng/ml (day 8) GM-CSF and 20ng/ml IL-4. Loosely adherent cells were harvested at day 10 and stimulated with 1μg/mL LPS (Sigma-Aldrich) in 37°C incubator for 24 hours. Cells were then pulsed with 5μg/ml of LCMV-GP_61-80_ peptides or left un-pulsed in bmDC culture medium for 2 hours at 37°C incubator. One million bmDCs were injected per mouse intravenously on the same day as tumor injection.

### *In vivo* antibody blockade, diphtheria toxin and tamoxifen treatments

For *in vivo* CTLA4 blockade, 300 µg/mouse of anti-CTLA4 (Clone UC10-4F10-11, BioXcell Cat# BE0032) or isotype (polyclonal Armenian Hamster IgG, clone: NA, BioXcell Cat#BE0091) antibodies were administered intravenously at day 0, 2 and 5 after tumor injection for early treatment and day 21, 23 and 26 for late treatment. For PDL1 blockade, 250µg antibodies (Clone 10 F.9G2, BioXcell Cat#BE0101) or isotype control (rat IgG2b Clone LTF-2, BioXcell, Cat#BE0090) were administrated i.v. at day 0, 2 and 5 following tumor injection. 500µg/mouse anti-CD25 (clone PC-61.5.3, BioXcell, Cat# BE0012,) in vivo depleting antibodies or isotype (Rat IgG1, clone HRPN, BioXcell Cat#BE0088) were administrated at day - 4 and -2, and depletion efficiency was confirmed with blood by Flow Cytometry. For CD28 agonist, 100μg/mouse anti-CD28 (clone PV-1, BioXcell, Cat#BE0015-5) or isotype (polyclonal Armenian hamster Armenian Hamster IgG, clone N/A, BioXcell, Cat#BE0091) antibodies were administrated at day 5 post tumor injection. For CD3 agonist, 100μg/mouse anti-CD3 (clone 145-2C11, BioXcell, Cat#BP0001-1) or isotype (polyclonal Armenian Hamster IgG, clone N/A, Cat#BP0091, BioXcell) antibodies were administrated at day 5 post tumor injection. For diphtheria toxin (DT)-mediated Treg depletion *in vivo*, 0.5µg DT was dissolved in 100ul PBS and administered retro-orbital at day 0 and 2 after tumor injection. Depletion of Tregs was confirmed via blood. For tamoxifen-mediated CTLA4 deletion, FoxP3-Cre x CTLA4 flox mice were treated daily for 5 days before tumor injection intraperitoneally with freshly made tamoxifen (2mg/mouse per treatment). Deletion of CTLA4 was confirmed by flow cytometric staining in blood.

### Flow cytometry and *ex vivo* peptide stimulation

Single-cell suspensions were prepared from organs and were stained *ex vivo* using antibodies in Table S3. Transcription factor stains were performed using FoxP3 Transcription Factor kit (eBiosciences, Cat No 00-5523-00). Samples were run on a FACS Verse or a FACS Lyric (BD Biosciences) and data analyzed using Flow Jo software (v.10; Treestar). For ex vivo peptide stimulation and cytokine staining, LN single cell suspension were stimulated for 5h at 37°C with 5ug/ml of LCMV-GP_61-80_ in the presence of 50 U/ml recombinant murine IL2 (Thermofisher P2747) and 1mg/ml Brefeldin A (Sigma). Cells were then stained with Zombie Aqua^TM^ Fixable Viability Kit (BioLegend), followed by extracellular staining. The cells were then washed, fixed, permeabilized (BioLegend cytokine staining kit) and stained with anti-cytokine antibodies.

### CRISPR-mediated CTLA4 deletion

CD4 SMARTA T cells were isolated from naïve mice and were washed and resuspended in 20 μl buffer P3 (P3 Primary Cell 4D-Nucleofector X Kit S, Lonza, cat. no. V4XP-3032). Between 10^5^ to 10^7^ cells were used per nucleofection. Single guide RNAs (sgRNAs) were incubated together with recombinant Cas9 (TrueCut Cas9 Protein v2, 5ug/ul, Thermo Fisher Scientific, cat. no. A36499) for 15 min at room temperature, at a ratio of 1:3.3 (i.e., 30 pmol Cas9 protein per 100 pmol sgRNA) to form the CRISPR–Cas9–sgRNA- ribonucleoprotein (RNP) complex. To increase deletion efficiency, a combination of three different pre-validated sgRNAs against CTLA-4 were mixed with the cell suspension (see list below), and transferred into a 16-well nucleocuvette strip (Lonza, cat. no. V4XP-3032). For negative control, sgRNA (TrueGuide™ sgRNA Negative Control, non-targeting 1; Thermo Fisher Scientific, cat. no. A35526t) was annealed with Cas9 at the same molar ratio. Cells were nucleofected using program DS137 and buffer P3 on the 4D- Nucleofector system (4D-Nucleofector X unit, Lonza, cat. no. AAF-1003X). Pre-warmed complete culture medium supplemented with IL-7 (10ng/ml) was added to each well and cells were transferred to a 96-well plate for recovery at 37°C for 6 hours before TAGIT label and transfer to mice that subsequently received tumor cells. The knockout efficiency based on CTLA-4 expression was confirmed by flow cytometry.

**sgRNA sequences against mouse CTLA-4 -** TrueGuide™ Synthetic sgRNA (Thermo Fisher Scientific) CRISPR58571_SGM

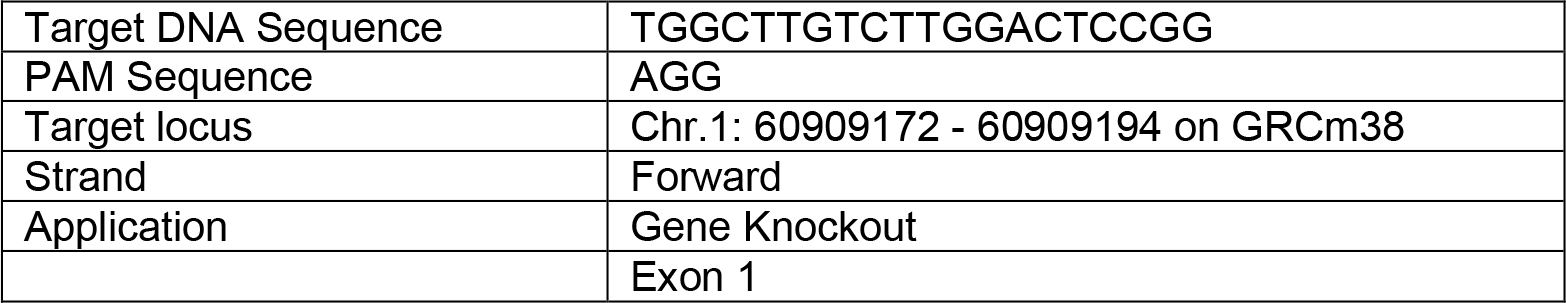

CRISPR58590_SGM

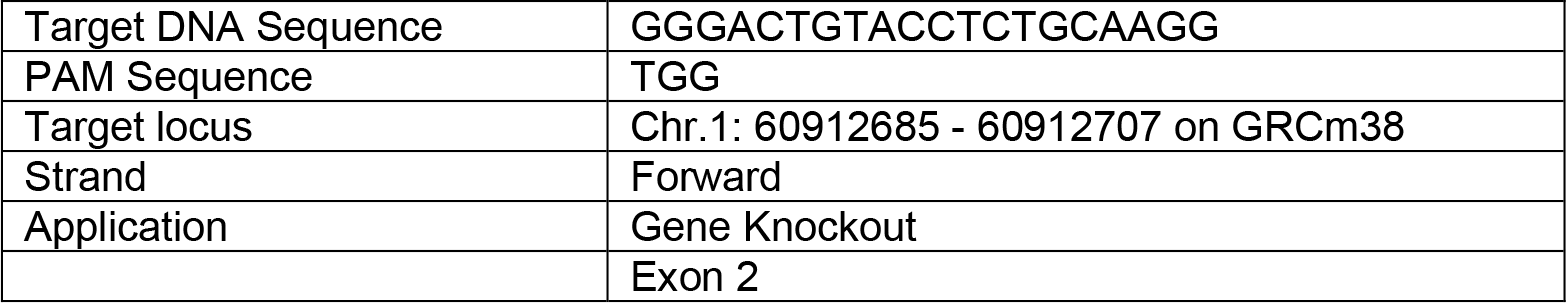

CRISPR58569_SGM

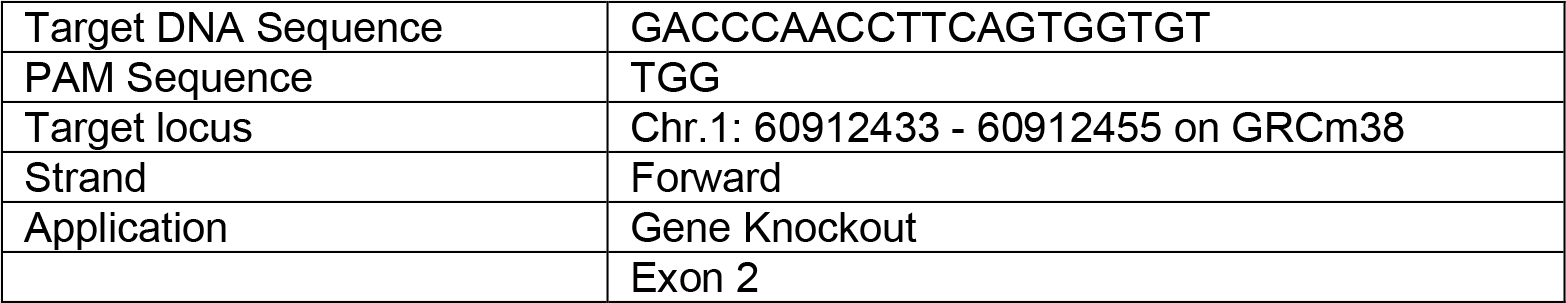

### Time-of-Flight mass cytometry (CyTOF)

CyTOF antibodies are listed in Table S1 and S2. Where indicated in Table S1, already conjugated antibodies were purchased from Fluidigm. Conjugation of purified antibodies was performed at the SickKids-UHN Flow and Mass Cytometry Facility using the MaxPar Antibody Labeling Kit (Fluidigm). All antibodies were titrated prior to the experiments. CD4 T cells were enriched with autoMACS Pro Separator (Miltenyi Biotec) using CD4 (L3T4) microbeads (order no. 130-117-043, Miltenyi Biotec).

For T cell phenotype panel: Samples were washed with PBS and stained with 12.5μM cisplatin (Cedarlane/biovision 1550-1000) in PBS for 1 min at room temperature before quenching with FBS. Samples were stained for 15 minutes at room temperature with FcR blockade and with antibodies that did not stain well after fixation (labelled in red in Tables) . Cells were fixed and permeabilized for 10 minutes at room temperature with Transcription Factor Staining Kit (eBioscience, Cat No 00-5523-00), and then barcoded with Cell-ID 20-Plex Pd Barcoding Kit (Fluidigm, Product No 201060). Combined samples were then incubated with surface antibody cocktail (Table S1) for 30 minutes at 4°C. After surface staining, samples were permeabilized and stained with intracellular antibodies for 30 minutes at room temperature using Transcription Factor Staining Kit (eBioscience, Cat No 00-5523-00). For the DNA stain, cells were incubated overnight in PBS (Multicell) with 0.3% (wt/vol) saponin, 1.6% (vol/vol) paraformaldehyde (Polysciences Inc.), and 1nM iridium (Fluidigm). Samples were acquired on a Helios mass cytometer (Fluidigm) at Sick Kids – UHN Flow and Mass Cytometry Facility or Princess Margaret Cancer Centre. EQ Four Element Calibration Beads (Fluidigm) were used to normalize signal intensity over time on CyTOF software version 6.7. FCS files were manually debarcoded and analyzed using R as described below.

For metabolism panel: Cells were washed with PBS and samples were labelled with 103Rh viability dye (for live/dead cell discrimination) at 37°C for 15min, followed by washing with PBS. Samples were then fixed and barcoded as described above. Following Fc receptor blocking for 10 min at room temperate, the pooled barcoded sample was incubated with the first surface cocktail of CD36-FITC and CD98-APC for 30 min at 4°C followed by washing. The cells were then stained with a second surface cocktail (Table S2) for 30 min at 4°C. The sample was stained with intracellular antibody staining for 30 min at room temperature. The DNA stain was then performed as described above.

### Bioinformatic Analyses (CyTOF)

Data pre-processing and dimensionality reduction of CyTOF data: Preprocessing of files was performed using FlowJo^TM^ Software (v10) software. Samples were manually debarcoded and exported as separated FCS files. SMARTA cells were filtered by gating on DNA, singlets, live cells and CD45.1+ CD45.2- CD4+ MHCII- TCRβ+ cells, and raw signal events were then exported as matrices in csv format. Data was then analyzed in R (v 4.1.0). All events were included in dimensionality reduction. Marker expression values were arcsinh transformed using a custom co-factor for each marker before clustering. Phenograph and UMAP were performed using the R implementation of the “Rphenograph” package (v 0.99.1) by JinmiaoChen lab on github (*68*) (https://github.com/JinmiaoChenLab/Rphenograph) and package “umap” (v 0.2.7.0). Differential state and abundance analyses were performed by “diffcyt” package (*69*) (v 1.14.0).

*Metabolic pathway scoring*: First, the sum of arcsinh-transformed expression of enzymes were calculated on a single cell level for each metabolic pathway (markers used to evaluate each pathway are defined as following), and then divided by the number of enzymes used to evaluate each metabolic pathway:

Glycolysis: GLUT1, HK2, GAPDH, MCT4

Glutamine and amino acid: CD98, GLS, GOT2, GLUD1/2

OXPHOS: IDH1, CS, ATP5A, CytC

Lipid: CD36, CPT1A, ACADM, pACC

Statistical significance of each pathway was calculated at the single cell level in R using Wilcox test only comparing between CD4 T_TS_ cells to Th1 and Tfh.

### Single cell RNA-sequencing

SMARTA cells were transferred into WT or FoxP3-DTR mice one day prior to tumor injection. Mice were treated with DT or PBS at day 0 and 2 post PyMG tumor injection. Anti-CTLA4 antibody was injected at day 0, 2 and 5 after tumor injection. Mice were sacrificed at day 8 and inguinal lymph nodes were harvested and digested into single cell suspension. CD4 T cells were enriched on autoMACS Pro Separator (Miltenyi Biotec) with CD4 positive selection beads (clone L3T4, Order no. 130-117-043, Miltenyi Biotec), followed by FACSorted on a Moflo Astrios (Beckman Coulter) or a BD FACSaria Fusion cell sorter for SMARTA cells based on CD45.1 expression. After sorting and purity check on flow cytometer, SMARTA cells were then resuspend in 10X Genomics Chromium single cell RNA master mix and loaded onto a 10x chromium chip. cDNA Library was generated using the Chromium Single Cell 3’ Reagents Kits v2 User Guide:CG00052 Rev B, and then sequenced on the Illumina HiSeq 2500 platform to achieve an average of 40,000 reads per cell. Sequencing was performed at the Princess Margaret Genomics Center. TAGIT expression on SMARTA cells were checked before and after sorts to make sure the procedures did not preferentially kill the effector population.

### Bioinformatic Analyses (mouse scRNA-Seq)

#### Reads alignment and quantification

Base calling was performed using Illumina RTA (v 3.4.4) and bcl2fastq2 (v 2.20) to generate bcl files. Cell ranger (v 6.0.0) was then used to demultiplex bcl into fastq files and to align reads to the mm10 genome (refdata-gex-mm10-2020-A downloaded from the 10x genomics website).

#### Data preprocessing

*Quality control and normalization:* Data was further analysed using R (v 4.1.1) and the R Seurat (v 4.0.4) package (Hao et al. 2021). Cells meeting the below criteria were considered as good quality cells and were kept for subsequent analyses:

- Naïve: 500 < genes expressed < 2300 and mitochondrial genes (%) < 10 (7955 cells)
- Isotype: 500 < genes expressed < 6000 and mitochondrial genes (%) < 12 (1099 cells)
- aCTLA4: 1000 < genes expressed < 6000 and mitochondrial genes (%) < 7 (4566 cells)
- Treg depletion: 700 < genes expressed < 7000 and mitochondrial genes (%) < 12 (2186 cells)
- Treg depletion + aCTLA4: 600 < genes expressed < 6000 and mitochondrial genes (%) < 13 (4343 cells).

Genes expressed in less than 3 cells and more than 12,000 cells were excluded from further analyses. Gene expression of all samples were initially normalized using the LogNormalize method and adjusted to regress out percentage of mitochondrial genes and cell cycle genes. Then data was exported to the SeqGeq application (v 1.8) where raw counts were normalized using the scTransform normalization and clustered using Louvain algorithm and the resolution indicated in the respective figure legends using the Seurat plug-in (4.0.4) (*70*) of SeqGeq.

#### Gene set variation analysis (GSVA)

We estimated activity of Hallmark gene sets taken from MSigDB (PMID: 23323831 and 21546393) on cellular level using the GSVA (v 1.40.0) and msigdbr (v 7.4.1) R packages and the R project (v 4.1.1) with the following parameters: kcdf=“Poisson”, method=’gsva’, min.sz=5, max.sz=500, parallel.sz=13. Enrichment scores for each cluster were then calculated as the average gsva scores of cells in each cluster. A two-sided Mann Whitney U test was performed to test whether gsva scores of celles in each cluster are significantly different than those in cells belonging to the remaining clusters. P-values were then corrected for multiple hypotheses testing using the FDR method. Gene sets having an FDR < 0.05 for at least one cluster were retained.

#### Monocle trajectory analysis

To determine cell trajectory, we used monocle3 (v 1.0.0) R package with R (v 4.1.1) (PMID: 24658644). Seurat clusters were used for this analysis. Graph was learnt using learn_graph_control=list(ncenter=170) for the isotype sample and ncenter=260 for all samples. Cells were ordered and pseudotime was calculated using default parameters.

#### Venn Diagrams

First, 6 differential expression analyses were performed between (1) paralyzed CD4 T_TS_ cells vs naïve CD4 T cells, (2) effector virus-specific CD4 T cells vs naïve CD4 T cells, (3) exhausted virus-specific CD4 T cells vs naïve CD4 T cells, (4) paralyzed CD4 T_TS_ cells vs naïve CD8 T cells, (5) effector virus-specific CD8 T cells vs naïve CD8 T cells, (6) exhausted virus-specific CD8 T cells vs naïve CD8 T cells. The top 500 up-regulated genes of the first 3 analyses make the left panel and the top 500 up- regulated genes of the last 3 analyses make the right panel in Figure 4E. Each dataset was first compared to naïve cells to identify differentially expressed genes (DEGs) unique to each T cell population, separate from “general” T cell expressed genes. The different data sets were procured from: scRNA-seq data for effector virus-specific CD4 T cells (*51*); exhausted virus-specific CD4 T cells (*52*); naive virus-specific CD8 T cells (*50*); effector virus-specific CD8 T cells (*50*) and exhausted virus-specific CD8 T cells (*50*).

#### GSEA and IPA

*Gene Set Enrichment Analysis* (*71, 72*) was performed by GSEA application (Broad Institute v. 4.2.2) using the Enrichment Map gene sets from ImmuneSigDB “GSE14308_TH17_VS_NAIVE_CD4_TCELL_UP” (*73*) (for Th17), “GSE14308_TH2_VS_NAIVE_CD4_TCELL_UP” (*73*) (for Th2), “GSE14415_INDUCED_TREG_VS_TCONV_UP” (*74*) (for iTreg), and Th1 and Tfh gene sets generated in previous Brooks Lab project(*9*). Gene lists were generated and exported from SeqGeq and preranked based on P value in R using the formula sign(log(gene.list$FoldChange))*(-log10(gene.list$Pvalue)). Ingenuity Pathway Analysis software (Qiagen) was used for the predicted pathway and upstream regulator analysis based on the gene lists generated from SeqGeq.

### Human scRNA-Seq data analyses

scRNA-Seq data was taken from two previously published studies of breast (*53*) (GEO: GSE180286) and lung adenocarcinoma (*54*) (GEO: GSE131907) cancers; data was reanalyzed from scratch as described below.

Breast cancer: samples were obtained from lymph nodes of 5 breast cancer patients (2 biological replicates from each patient), one of these samples (P3_LN1) was considered an outlier, as it has, on average, more expressed genes, read counts and mitochondrial genes percentage per cell than all other samples, so it was excluded from further analyses. Lung cancer: samples were obtained from lymph nodes of 17 lung cancer patients.

We filtered out dead and poor-quality cells and doublets by excluding cells meeting the following criteria: nFeature_count < 200, nFeature_count > nFeature_count_cutoff, mitochondrial genes (%) > mitochondrial_percentage_cutoff for each sample. Cutoffs were chosen in a way to remove the cells at the upper tail of the nFeature_count and mitochondrial_percentage distributions. Used cutoffs and number of cells before and after filtering can be found in Table S4.

We used Seurat (v 4.3.0) and R (v 4.2.2) to merge and logNormalize samples of each cancer type apart. All samples were then integrated using the STACAS (v 2.0.2) algorithm using default parameters (PubmedID: 32845323). After integration of all samples together, we scaled data, ran UMAP using the first 26 principal components, ran FindNeighbors and ran FindClusters using resolution 0.3. Next, T cells were identified based on CD3E and TRAC expression profiles as cells in clusters: 0, 1, 3 and 6. Data was then re-clustered with resolution 0.8, and CD4 T cells. Subsequently, we re-clustered cells using resolution 0.8 and identified CD4 T cells by annotating cells using SingleR (2.0.0) (used parameters: de.method=“wilcox”, sd.thresh=1.2) and Monaco (GSE107011) as a reference data set (Pubmed ID: 30726743) on cluster level (resolution 0.8) and obtained 55,521 CD4 T cells. To identify paralyzed CD4 T cells in human, we considered the 373 genes in Figure 4G (left panel, genes in blue circle) as a signature of the tumor-specific CD4 paralyzed in mouse and used their orthologs in human to identify similar cells in human. To this end, we used the cosine similarity applied on 1) mean expression of the 373 genes in mouse and their human orthologs in each cell of the 55,521 cells in human. Mouse to human orthologs mapping was taken from MGI reports (http://www.informatics.jax.org/downloads/reports/). Human paralyzed CD4 T cells were identified by cells having a z-score of cosine similarity > 2.5, resulting in 616 cells. We then defined a signature of human paralyzed CD4 T cells as the average expression of the top 25 up-regulated genes in human paralyzed CD4 T cells to all other CD4 T cells in human samples using the findMarkers function of Seurat. The Signature Expression was quantified in each cell as the Sum(normalized expression of gene1, gene2…gene25)/25. Finally, we performed biological processes GO enrichment analysis using gProfileR (PubmedID: 31066453) with default parameters, except that we treated the list of 25 genes as an ‘ordered query’.

### Statistical analyses (other than CyTOF and scRNAseq)

Mann-Whitney U test or two-way ANOVA were calculated by GraphPad Prism 9 (GraphPad Software, Inc.).

## Acknowledgements

We thank past and present members of the Brooks laboratory for technical help and discussion. This work was supported by the Canadian Institutes of Health Research (CIHR) Foundation Grant FDN148386 (D.G.B), the National Institutes of Health (NIH) grant AI085043 (D.G.B), the Scotiabank Research Chair to D.G.B.

## SUPPLEMENTAL FIGURES

**Figure S1. CD4 T cell help is important for long-term tumor control, yet natural CD4 T cell response remains suboptimal. Related to Figure 1.**

(A) Tumor size of B16-F10 in WT and CD4 knockout (KO) mice at the indicated time points.

(B) Growth kinetics of MC38 and PyMT in WT or mice depleted of Tregs at the onset of tumor implantation.

(C) Growth kinetics of PyMG versus PyMT (left), and MC38GP versus MC38 (right).

(D) WT mice receiving either empty or GP_61-80_ peptide-pulsed β2m-/- bmDCs same day of PyMG tumor transplantation. Box plot shows the tumor size at the indicated time points.

(E) Mice received CD4 SMARTA T_TS_ cells and PyMG tumors, followed by either unlabeled or GP_61-80_ peptide labeled β2m-/- bmDC. Flow plots and bar graph show the frequency of SMARTA cells out of total PyMG tumor-infiltrating CD4 T cells at day 8 or day 30.

(F) Mice received PyMG. 20 days later, the mice received 1 million β2m-/- bmDC labeled with the LCMV- GP67-80 peptide (red) or irrelevant peptide (the OVA peptide specific for OT2 cells; blue). Tumor growth was measured longitudinally.

Data represents 3 independent experiments with at least 5 mice per group. Error bars indicate standard deviation (SD). For tumor growth kinetics, significance is determined by two-way ANOVA; other panels, significance is determined by Mann-Whitney U test. *, p<0.05.

**Figure S2. Characteristics of CD4 T_TS_ cell paralysis is not observed in CD8 T cells. Related to Figure 1.**

(A-D) TAGIT-labelled naïve SMARTA cells were transferred to WT mice followed by PyMG injection.

(A) Percentage of proliferated SMARTA T_TS_ cells in dLN at each time point after tumor initiation.

(B) Expression of CD44 and CD86 versus dilution of proliferation dye by SMARTA T_TS_ cells on day 8 dLN.

(C) Proliferation of tumor-specific CD4 SMARTA T cells in the tumor at day 8.

(D) Number of SMARTA cells in each proliferation peak from Figure 1C in the dLN at the indicated day after PyMG administration.

(E) Naïve TAGIT-labeled CD4 SMARTA T cells or ovalbumin-specific CD4 OT-II T cells were transferred to WT mice followed by injection of MC38-GP or MC38-expressing ovalbumin (MC38-OVA), respectively. Histograms show TAGIT dilution and bar graphs indicate the number of SMARTA or OT-II cells in the dLN.

(F) Proliferation (histograms) and total number (bar graph) of CD8 P14 T_TS_ cells in dLN at each time point after PyMG initiation.

(G) Percentage (flow plots) of CD8 P14 T_TS_ cells in the tumor (gated on total CD8 TILs). Number (bar graph) of CD8 P14 T_TS_ cells in the tumor.

(H) TAGIT-labeled naïve SMARTA cells were transferred to WT mice 21 days after PyMG implantation. TAGIT dilution (histograms) and percentage of proliferation (bar graph) on day 29 and 36 of tumor initiation (day 8 and 15 respectively after SMARTA T cell transfer)

Data represents 3 independent experiments with at least 5 mice per group. Error bars indicate SD. Significance determined by Mann-Whitney U test. *, p<0.05.

**Figure S3. Tumor-specific CD4 T cell differentiation. Related to Figure 2.**

(A) Representative expression of RORγt and FoxP3 on SMARTA T_TS_ cells in dLN at each time points.

(B) TCF1 and Bcl6 expression by naïve and tumor-activated (day 8) SMARTA T_TS_ cells in dLN.

(C) Number of SMARTA cells from dLN expressing TNFα or IL2 after ex vivo LCMV-GP_61-80_ peptide stimulation.

(D) IFNγ and IL10 expression by SMARTA T_TS_ cells after ex vivo LCMV-GP_61-80_ peptide stimulation (day 8).

(E) (Left) Bar graph shows the frequency of each cluster from the UMAP in Figure 2E. (Right) Differential abundance (calculated by diffcyt, colored based on log2 fold-change, *adjusted p<0.05) comparing the frequency of each cluster on day 8 to day 15, and day 15 to day 30. Red is increasing with time, blue is decreasing with time, and grey means the changes are not significant.

(F) Differential states analysis (calculated by diffcyt) of the change in expression of the indicated protein in each cluster from day 8 to 30. Red equals increased at day 30 and blue decreased at day 30. *, adjusted p-value<0.05 (from Figure 2E).

(G) Expression of SLAMF1 and Tbet protein by naïve c9 (blue) and iTreg c8 (red) CD4 SMARTA T_TS_ cells (from Figure 2E).

(H) Expression of Blimp1 and Bcl6 by the indicated clusters (from Figure 2E).

Data represents 2-3 independent experiments with at least 4 mice per group. Error bars indicate SD. Significance determined by Mann-Whitney U test. *, p<0.05.

**Figure S4. Metabolic CyTOF and scRNA-seq analysis of CD4 T_TS_ cells. Related to Figure 3 and Figure 4.**

(A) Violin plots indicate the single cell arcsinh-transformed intensity of the indicated protein. Statistics were analyzed via diffcyt as indicated in Figure 3C.

(B) Violin plot depicting the single-cell pseudotime differentiation scores of each cluster. The pseudotime starting point is in the naïve-like c3 from Figure 4C.

(C) Heatmap depicts the complete result of GSVA pathway analysis from Figure 4D. Color depicts the z-score of the mean GSVA single cell scores of activation level of each pathway. P-value is calculated by Mann Whitney U test, ***, p<0.001; **, p<0.01; *, p<0.05. Pathways in red are the same pathways in Figure 4D.

(D) UMAP plots show the Seurat clustering (resolution 0.3) of naïve (left) and tumor-activated (right) SMARTA T_TS_ cells. The bar graph shows the frequency of the clusters within each sample. (E) IPA-predicted upstream regulator of cells in cluster 4 (from Figure S4D). All regulator predictions are significant with p<0.05. Red words indicate regulators discussed in the results.

(F) Box plot shows the percentage of human CD4 T cells sharing the mouse-derived T_TS_ cell signature (of total sequenced CD4 T cells) in the dLN samples.

(G) Heatmap depicts the expression level of top 25 differentially expressed genes (DEGs) in human CD4 T cells compared to the other CD4 T cells in the lymph nodes from breast (top) and lung (bottom) cancer patients. Box plots shows the single cell expression of the combined T_TS_ signature. Red box represents cells with the CD4 T_TS_ signature and blue box indicates other CD4 T cells in breast (top) and lung (bottom) cancer. The box represents the mean to the 75^th^ and 25^th^ quartile. *, p<2.2×10^-16^

(H) GO term pathway enrichment analysis of the top 25 up-regulated genes in human CD4 T cells with the mouse-derived T_TS_ signature vs all other cells.

Except for human data, data are representative of 2 experiments with at least 3 mice per group.

**Figure S5. Impact of CTLA4 blockade on CD4 T_TS_ cell paralysis. Related to Figure 5.**

(A) Proliferation of CD4 SMARTA T_TS_ cells in dLN of MC38GP tumor following isotype or anti-PDL1 antibody treatment.

(B-D) On day 0, 2 and 5 after PyMG administration, mice were treated with CTLA4 blocking or isotype control antibodies and CD4 T cells in dLN stained by CyTOF on day 8.

(B) UMAP plots overlaid with PhenoGraph clusters of SMARTA T_TS_ cells (samples were concatenated within each group for the UMAP presentation). The Bar graph shows the frequency of each cluster, with circles in each bar representing individual mice. *: adjusted p<0.05, diffcyt differential abundance test.

(C) Heatmap depicts normalized z-scores of the arcsinh-transformed MSI of the indicated protein for each cluster.

(D) Heatmap depicts the differential state analysis of protein expression by each cluster comparing anti- CTLA4-treated to isotype group (*=FDR<0.05), and coloration presents log2 fold-change.

(E) FoxP3-iCRE x CTLA4 fl/fl mice were treated with vehicle control (blue) or tamoxifen (red). First histogram (left) shows CTLA4 expression on FoxP3+ Tregs in the blood prior to SMARTA cell transfer and PyMG administration. Second histogram (right) shows the proliferation at day 8 in dLN in vehicle control (blue) or tamoxifen (red) treated FoxP3-iCRE x CTLA4 fl/fl mice. The bar graph shows the percentage of proliferation.

(F) CRISPR-mediated deletion of CTLA4 in naïve SMARTA T_TS_ cells. CTLA4 expression on and proliferation of CTLA4-deleted or control SMARTA T_TS_ cells in dLN at day 8 after PyMG initiation. Tregs are included for comparison of CTLA4 expression.

Data represents 2-3 independent experiments with at least 4 mice per group. Error bars indicate SD. Significance determined by Mann-Whitney U test, except for CyTOF analyses.

**Figure S6. Impact of Treg depletion on CD4 T_TS_ cell paralysis. Related to Figure 5.**

(A) The frequency of FoxP3+ Treg cells (non-SMARTA cells) of total CD4 T cells in the dLN after PyMG initiation.

(B) Number of SMARTA T_TS_ cells in day 8 dLN of FoxP3-DTR mice treated with PBS or DT.

(C) Mice were treated twice (day -4 and day -2) prior to PyMG injection with either 500µg CD25 depleting or isotype antibodies. Blood was stained to confirm depletion of CD25+ cells prior to SMARTA cell transfer and tumor injection. Histograms show the proliferation of SMARTA cells in dLN on day 8. The bar graph shows the number of SMARTA cells in dLN.

(D) FoxP3-DTR mice received SMARTA cells followed a day later by administration of PyMG tumor cells. On day 21 and 23 mice were treated with PBS or DT. Bar graph depicts the total number of SMARTA T_TS_ cells in dLN at day 29 (8 days after initial PBS or DT treatment).

(E-H) FoxP3-DTR mice received TAGIT-labeled naïve SMARTA cells followed by PyMG injection. On day 0 and 2 the mice were treated with either PBS or DT and CyTOF was performed in the dLN on day 8.

(E) Arcsinh-scaled expression of the indicated protein by SMARTA T_TS_ cells.

(F) Histogram shows the FoxP3 expression by CD4 SMARTA T_TS_ cells from dLN of isotype control (blue), Treg-depleted (yellow) and aCTLA4-treated (red) mice

(G) Arcsinh-scale expression of the indicated protein by SMARTA T_TS_ cells.

(H) Expression of the indicated protein by CD4 SMARTA iTreg (c5, 7 and 8 from Figure 5D) and Tregs (FoxP3+ Helios+ of non-SMARTA CD4 T cells in PBS dLNs). Bar graphs show the percentage and GMSI of the indicated protein by SMARTA iTreg (red) and non-SMARTA Tregs (blue).

(I-J) (I) DC analysis in WT C57Bl/6 mice treated with isotype control or anti-CTLA4 blocking antibody on day 0, 2 and 5 after PyMG initiation. (J)DC analysis in DEREG mice treated with PBS as control or diphtheria toxin to deplete Tregs on day 0 and 2 after PyMG initiation. DC were gated based on high CD11c and MHC II expression (left plots in each group). Two subsets of CD11c MHC II high DC were identified. These were then further defined based on CD8a and CD11b expression as DC1 (CD8 T cell stimulating) and two populations of DC2 (CD4 T cell stimulating) (right plots in each group). Bar graphs show the number of DC in each subset, and their expression of MHC II, CD80 and CD86 (expression data is from CyTOF and is shown as geometric mean signal intensity).

(K) Following PyMG administration, FoxP3-DTR mice were treated with PBS or DT; and CTLA4-blocking or isotype antibodies. The bar graphs show the frequency of IL2 and TNFα producing CD4 SMARTA T cells after *ex vivo* LCMV-GP_61-80_ peptide stimulation.

Data represents 2-3 independent experiments with at least 4 mice per group. Error bars indicate SD. Significance determined by Mann-Whitney U test.

**Figure S7. Altered transcriptional regulation of CD4 T_TS_ cells following single or combined Treg depletion and CTLA4 blockade. Related to Figure 6.**

(A) Top 10 DEGs (heatmap) and *Foxp3* and *Ikzf2* expression (UMAPs) of each SMARTA T_TS_ cell cluster from the scRNA-seq analysis.

(B) Upstream regulator activities determined by IPA (red = predicted increased activity, blue = predicted decreased activity). All shown are significant with p<0.05. Red indicates regulators discussed in results.

(C) IPA Pathway analysis Pathways activated or inhibited by IPA analysis. All pathways are significant with p<0.05.

**Figure S8. Model of CD4 T_TS_ cell paralysis and restoration.**

(I) Tregs reduce DC activation level in dLN by lowering surface MHCII and B7 family molecules (CD80 and CD86). Concurrently, activation-induced CTLA4 on CD4 T_TS_ cells further inhibits T cell priming by directly interfering with TCR signals and competing with CD28 for binding with CD80 and CD86. This integrated, two-tiered mechanism of Treg and CTLA4 mediated suppression results in subdued T cell activation, leading to the paralyzed state.

(II) Removing Tregs from dLN alleviates the suppression on DCs, increases MHCII and B7 family molecules, which restores proper T cell priming and proliferation. Despite the enhanced proliferation, CTLA4 expression by CD4 T_TS_ direct their differentiation towards highly active tumor specific iTregs.

(III) Blocking CTLA4 signals concurrent with Treg depletion potentiates effector differentiation of effector tumor-specific T helper cells and the production of IFNγ.

DC (dendritic cell); dLN (draining lymph node); Treg (natural Tregs); iTregs (induced Tregs).

## REFERENCES

1. A. D. Waldman, J. M. Fritz, M. J. Lenardo, A guide to cancer immunotherapy: from T cell basic science to clinical practice. Nature Reviews Immunology 20, 651–668 (2020).

2. K. E. Pauken, E. J. Wherry, Overcoming T cell exhaustion in infection and cancer. Trends Immunol 36, 265–276 (2015).

3. N. N. Hunder et al., Treatment of metastatic melanoma with autologous CD4+ T cells against NY-ESO-1. N Engl J Med 358, 2698–2703 (2008).

4. E. Tran et al., Cancer immunotherapy based on mutation-specific CD4+ T cells in a patient with epithelial cancer. Science 344, 641–645 (2014).

5. T. Ahrends et al., CD4+ T Cell Help Confers a Cytotoxic T Cell Effector Program Including Coinhibitory Receptor Downregulation and Increased Tissue Invasiveness. Immunity 47, 848–861.e845 (2017).

6. C. Cui et al., Neoantigen-driven B cell and CD4 T follicular helper cell collaboration promotes anti-tumor CD8 T cell responses. Cell 184, 6101–6118.e6113 (2021).

7. S. C. Wei et al., Distinct Cellular Mechanisms Underlie Anti-CTLA-4 and Anti-PD-1 Checkpoint Blockade. Cell 170, 1120–1133.e1117 (2017).

8. D. G. Brooks, L. Teyton, M. B. Oldstone, D. B. McGavern, Intrinsic functional dysregulation of CD4 T cells occurs rapidly following persistent viral infection. J Virol 79, 10514–10527 (2005).

9. L. M. Snell et al., Dynamic CD4+ T cell heterogeneity defines subset-specific suppression and PD-L1-blockade-driven functional restoration in chronic infection. Nature Immunology 22, 1524–1537 (2021).

10. A. Oxenius et al., Variable fate of virus-specific CD4(+) T cells during primary HIV-1 infection. Eur J Immunol 31, 3782–3788 (2001).

11. L. M. Fahey et al., Viral persistence redirects CD4 T cell differentiation toward T follicular helper cells. J Exp Med 208, 987–999 (2011).

12. T. Duhen et al., Co-expression of CD39 and CD103 identifies tumor-reactive CD8 T cells in human solid tumors. Nature Communications 9, 2724 (2018).

13. A. Reuben et al., Comprehensive T cell repertoire characterization of non-small cell lung cancer. Nature Communications 11, 603 (2020).

14. P. C. Rosato et al., Virus-specific memory T cells populate tumors and can be repurposed for tumor immunotherapy. Nature Communications 10, 567 (2019).

15. W. Scheper et al., Low and variable tumor reactivity of the intratumoral TCR repertoire in human cancers. Nat Med 25, 89–94 (2019).

16. N. N. Hunder et al., Treatment of Metastatic Melanoma with Autologous CD4+ T Cells against NY-ESO-1. New England Journal of Medicine 358, 2698–2703 (2008).

17. E. Shklovskaya et al., Tumour-specific CD4 T cells eradicate melanoma via indirect recognition of tumour-derived antigen. Immunol Cell Biol 94, 593–603 (2016).

18. Y. Xie et al., Naive tumor-specific CD4(+) T cells differentiated in vivo eradicate established melanoma. J Exp Med 207, 651–667 (2010).

19. A. Schietinger et al., Tumor-Specific T Cell Dysfunction Is a Dynamic Antigen-Driven Differentiation Program Initiated Early during Tumorigenesis. Immunity 45, 389–401 (2016).

20. R. Zander et al., CD4(+) T Cell Help Is Required for the Formation of a Cytolytic CD8(+) T Cell Subset that Protects against Chronic Infection and Cancer. Immunity 51, 1028–1042 e1024 (2019).

21. R. A. van Lier et al., Tissue distribution and biochemical and functional properties of Tp55 (CD27), a novel T cell differentiation antigen. J Immunol 139, 1589–1596 (1987).

22. L. Cimmino et al., Blimp-1 attenuates Th1 differentiation by repression of ifng, tbx21, and bcl6 gene expression. J Immunol 181, 2338–2347 (2008).

23. E. L. Pearce, Metabolism in T cell activation and differentiation. Curr Opin Immunol 22, 314–320 (2010).

24. S. Kouidhi, A. B. Elgaaied, S. Chouaib, Impact of Metabolism on T-Cell Differentiation and Function and Cross Talk with Tumor Microenvironment. Frontiers in Immunology 8, (2017).

25. F. J. Hartmann et al., Single-cell metabolic profiling of human cytotoxic T cells. Nature Biotechnology 39, 186–197 (2021).

26. R. J. Johnston et al., Bcl6 and Blimp-1 are reciprocal and antagonistic regulators of T follicular helper cell differentiation. Science 325, 1006–1010 (2009).

27. C. H. Chang et al., Posttranscriptional control of T cell effector function by aerobic glycolysis. Cell 153, 1239–1251 (2013).

28. O. Minchenko, I. Opentanova, D. Minchenko, T. Ogura, H. Esumi, Hypoxia induces transcription of 6-phosphofructo-2-kinase/fructose-2,6-biphosphatase-4 gene via hypoxia-inducible factor-1alpha activation. FEBS Lett 576, 14–20 (2004).

29. H. Zhang et al., HIF-1α activates hypoxia-induced PFKFB4 expression in human bladder cancer cells. Biochem Biophys Res Commun 476, 146–152 (2016).

30. M. O. Johnson et al., Distinct Regulation of Th17 and Th1 Cell Differentiation by Glutaminase-Dependent Metabolism. Cell 175, 1780–1795.e1719 (2018).

31. M. Ahmadian et al., PPARγ signaling and metabolism: the good, the bad and the future. Nature Medicine 19, 557–566 (2013).

32. M. Hernandez-Quiles, M. F. Broekema, E. Kalkhoven, PPARgamma in Metabolism, Immunity, and Cancer: Unified and Diverse Mechanisms of Action. Front Endocrinol (Lausanne) 12, 624112 (2021).

33. G. Ye et al., PPARα and PPARγ activation attenuates total free fatty acid and triglyceride accumulation in macrophages via the inhibition of Fatp1 expression. Cell Death Dis 10, 39 (2019).

34. G. Chinetti et al., PPAR-alpha and PPAR-gamma activators induce cholesterol removal from human macrophage foam cells through stimulation of the ABCA1 pathway. Nat Med 7, 53–58 (2001).

35. S. B. Widenmaier et al., NRF1 Is an ER Membrane Sensor that Is Central to Cholesterol Homeostasis. Cell 171, 1094–1109.e1015 (2017).

36. J. Jain, P. G. McCaffrey, V. E. Valge-Archer, A. Rao, Nuclear factor of activated T cells contains Fos and Jun. Nature 356, 801–804 (1992).

37. R. J. Martinez et al., Arthritogenic self-reactive CD4+ T cells acquire an FR4hiCD73hi anergic state in the presence of Foxp3+ regulatory T cells. J Immunol 188, 170–181 (2012).

38. D. García-Bernal et al., RGS10 restricts upregulation by chemokines of T cell adhesion mediated by α4β1 and αLβ2 integrins. J Immunol 187, 1264–1272 (2011).

39. Y. Jiang et al., Deficiency of programmed cell death 4 affects the balance of T cell subsets in hyperlipidemic mice. Mol Immunol 112, 387–393 (2019).

40. H. Lingel et al., CTLA-4-mediated posttranslational modifications direct cytotoxic T-lymphocyte differentiation. Cell Death Differ 24, 1739–1749 (2017).

41. E. Corapi, G. Carrizo, D. Compagno, D. Laderach, Endogenous Galectin-1 in T Lymphocytes Regulates Anti-prostate Cancer Immunity. Frontiers in immunology 9, 2190–2190 (2018).

42. F. Cedeno-Laurent et al., Galectin-1 inhibits the viability, proliferation, and Th1 cytokine production of nonmalignant T cells in patients with leukemic cutaneous T-cell lymphoma. Blood 119, 3534–3538 (2012).

43. H. Javanmard Khameneh et al., The Inflammasome Adaptor ASC Intrinsically Limits CD4+ T-Cell Proliferation to Help Maintain Intestinal Homeostasis. Frontiers in Immunology 10, (2019).

44. E. Sebzda, Z. Zou, J. S. Lee, T. Wang, M. L. Kahn, Transcription factor KLF2 regulates the migration of naive T cells by restricting chemokine receptor expression patterns. Nature Immunology 9, 292–300 (2008).

45. M. A. Weinreich et al., KLF2 transcription-factor deficiency in T cells results in unrestrained cytokine production and upregulation of bystander chemokine receptors. Immunity 31, 122–130 (2009).

46. J. Cao et al., The single-cell transcriptional landscape of mammalian organogenesis. Nature 566, 496–502 (2019).

47. C. Trapnell et al., The dynamics and regulators of cell fate decisions are revealed by pseudotemporal ordering of single cells. Nature Biotechnology 32, 381–386 (2014).

48. X. Qiu et al., Reversed graph embedding resolves complex single-cell trajectories. Nature Methods 14, 979–982 (2017).

49. H. Chi, Regulation and function of mTOR signalling in T cell fate decisions. Nature Reviews Immunology 12, 325–338 (2012).

50. Z. Chen et al., TCF-1-Centered Transcriptional Network Drives an Effector versus Exhausted CD8 T Cell-Fate Decision. Immunity 51, 840–855.e845 (2019).

51. A. Khatun et al., Single-cell lineage mapping of a diverse virus-specific naive CD4 T cell repertoire. J Exp Med 218, (2021).

52. R. Zander, A. Khatun, M. Y. Kasmani, Y. Chen, W. Cui, Delineating the transcriptional landscape and clonal diversity of virus-specific CD4+ T cells during chronic viral infection. eLife 11, e80079 (2022).

53. K. Xu et al., Single-cell RNA sequencing reveals cell heterogeneity and transcriptome profile of breast cancer lymph node metastasis. Oncogenesis 10, 66 (2021).

54. N. Kim et al., Single-cell RNA sequencing demonstrates the molecular and cellular reprogramming of metastatic lung adenocarcinoma. Nature Communications 11, 2285 (2020).

55. S. S. Ng et al., The NK cell granule protein NKG7 regulates cytotoxic granule exocytosis and inflammation. Nature Immunology 21, 1205–1218 (2020).

56. D. Bruniquel, N. Borie, S. Hannier, F. Triebel, Regulation of expression of the human lymphocyte activation gene-3 (LAG-3) molecule, a ligand for MHC class II. Immunogenetics 48, 116–124 (1998).

57. F. Annunziato et al., Opposite role for interleukin-4 and interferon-gamma on CD30 and lymphocyte activation gene-3 (LAG-3) expression by activated naive T cells. Eur J Immunol 27, 2239–2244 (1997).

58. L. M. McLane et al., Role of nuclear localization in the regulation and function of T-bet and Eomes in exhausted CD8 T cells. Cell Rep 35, 109120 (2021).

59. W. Scheper et al., Low and variable tumor reactivity of the intratumoral TCR repertoire in human cancers. Nature Medicine 25, 89–94 (2019).

60. K. E. Yost et al., Clonal replacement of tumor-specific T cells following PD-1 blockade. Nature Medicine 25, 1251–1259 (2019).

61. D. G. Brooks, D. B. McGavern, M. B. Oldstone, Reprogramming of antiviral T cells prevents inactivation and restores T cell activity during persistent viral infection. J Clin Invest 116, 1675–1685 (2006).

62. A. V. Gett, P. D. Hodgkin, A cellular calculus for signal integration by T cells. Nat Immunol 1, 239–244 (2000).

63. Y. Wu et al., FOXP3 controls regulatory T cell function through cooperation with NFAT. Cell 126, 375–387 (2006).

64. R. K. Naviaux, E. Costanzi, M. Haas, I. M. Verma, The pCL vector system: rapid production of helper-free, high-titer, recombinant retroviruses. J Virol 70, 5701–5705 (1996).

65. E. Kowarz, D. Löscher, R. Marschalek, Optimized Sleeping Beauty transposons rapidly generate stable transgenic cell lines. Biotechnol J 10, 647–653 (2015).

66. L. Mátés et al., Molecular evolution of a novel hyperactive Sleeping Beauty transposase enables robust stable gene transfer in vertebrates. Nat Genet 41, 753–761 (2009).

67. W. Xu et al., Early innate and adaptive immune perturbations determine long-term severity of chronic virus and Mycobacterium tuberculosis coinfection. Immunity 54, 526–541 e527 (2021).

68. J. H. Levine et al., Data-Driven Phenotypic Dissection of AML Reveals Progenitor-like Cells that Correlate with Prognosis. Cell 162, 184–197 (2015).

69. L. M. Weber, M. Nowicka, C. Soneson, M. D. Robinson, diffcyt: Differential discovery in high-dimensional cytometry via high-resolution clustering. Communications Biology 2, 183 (2019).

70. Y. Hao et al., Integrated analysis of multimodal single-cell data. Cell 184, 3573–3587.e3529 (2021).

71. A. Subramanian et al., Gene set enrichment analysis: A knowledge-based approach for interpreting genome-wide expression profiles. Proceedings of the National Academy of Sciences 102, 15545–15550 (2005).

72. V. K. Mootha et al., PGC-1α-responsive genes involved in oxidative phosphorylation are coordinately downregulated in human diabetes. Nature Genetics 34, 267–273 (2003).

73. G. Wei et al., Global mapping of H3K4me3 and H3K27me3 reveals specificity and plasticity in lineage fate determination of differentiating CD4+ T cells. Immunity 30, 155–167 (2009).

74. D. Haribhai et al., A central role for induced regulatory T cells in tolerance induction in experimental colitis. J Immunol 182, 3461–3468 (2009).

