## Supplementary Figures for "Molecular, metabolic and functional CD4 T cell paralysis impedes tumor control"

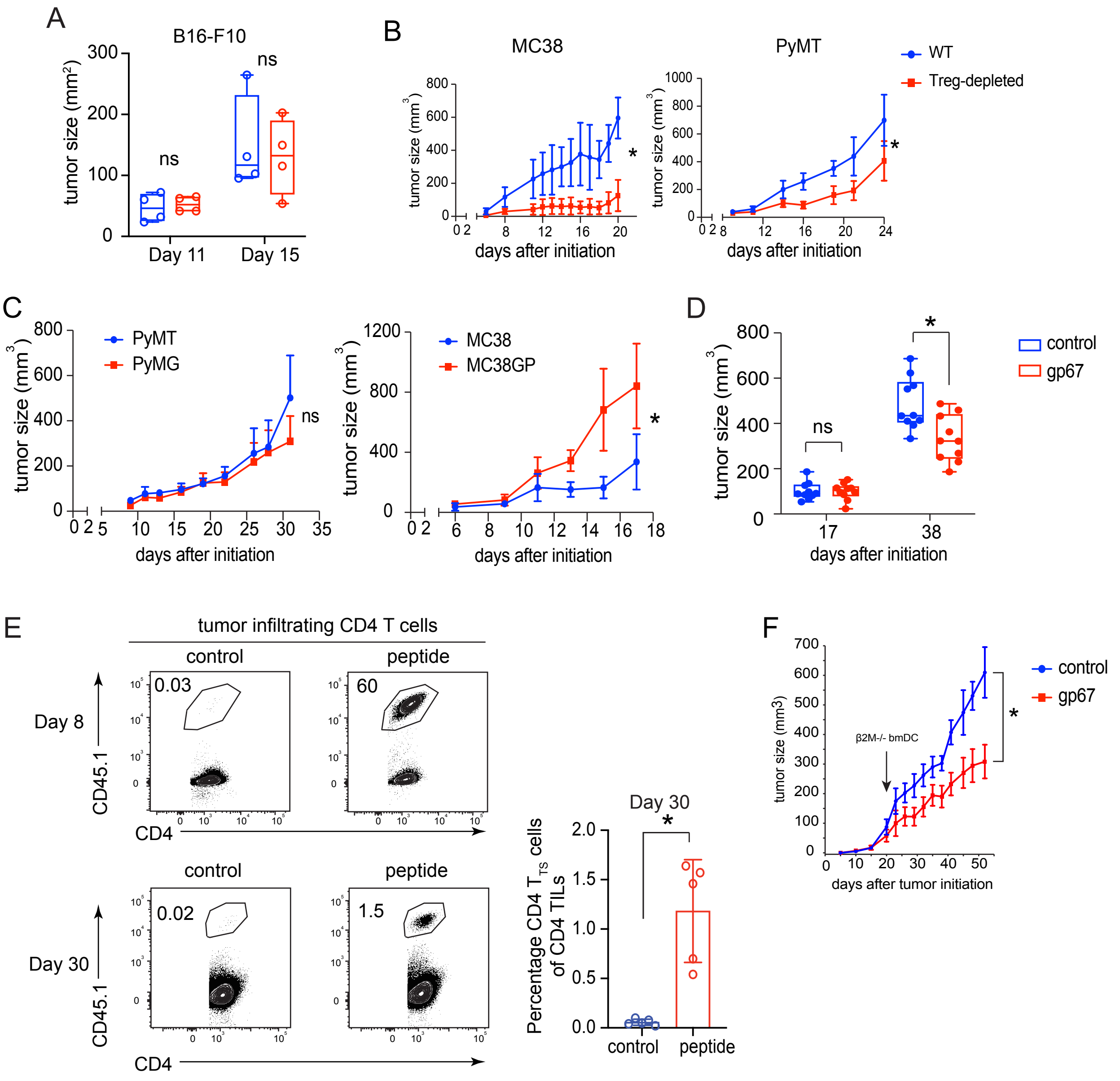
A
B16-F10
tumor size (mm2)
ns
Day 11
Day 15
B
MC38
PyMT
WT
Treg-depleted
tumor size (mm3)
days after initiation
C
PyMT
PyMG
MC38
MC38GP
D
control
gp67
E
tumor infiltrating CD4 T cells
control
peptide
Day 8
Day 30
CD45.1
CD4
0.03
60
0.02
1.5
Percentage CD4 TTS cells of CD4 TILs
F
β2M-/- bmDC
tumor size (mm3)
days after tumor initiation

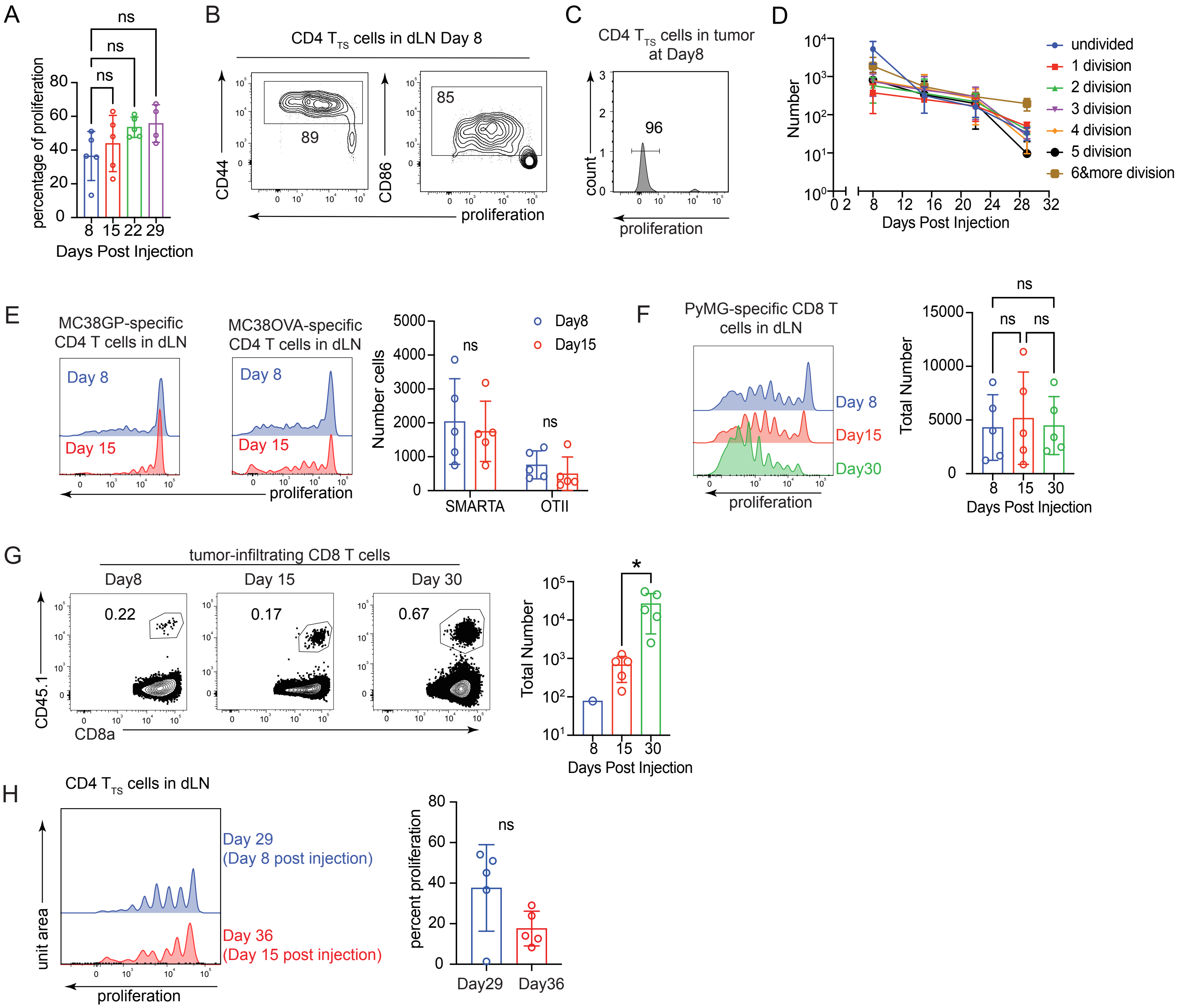


Supplementary Figure 2

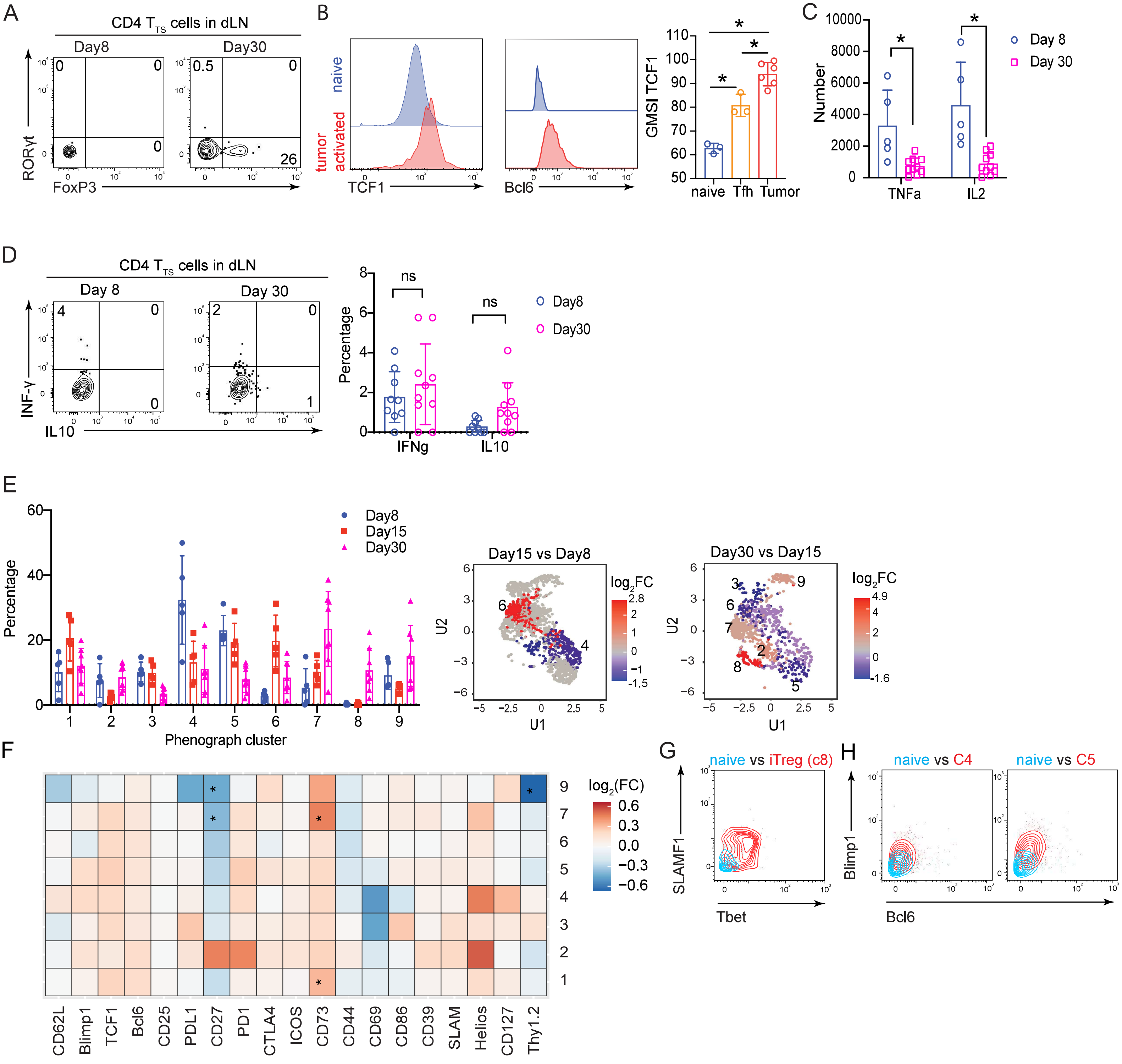


Supplementary Figure 3

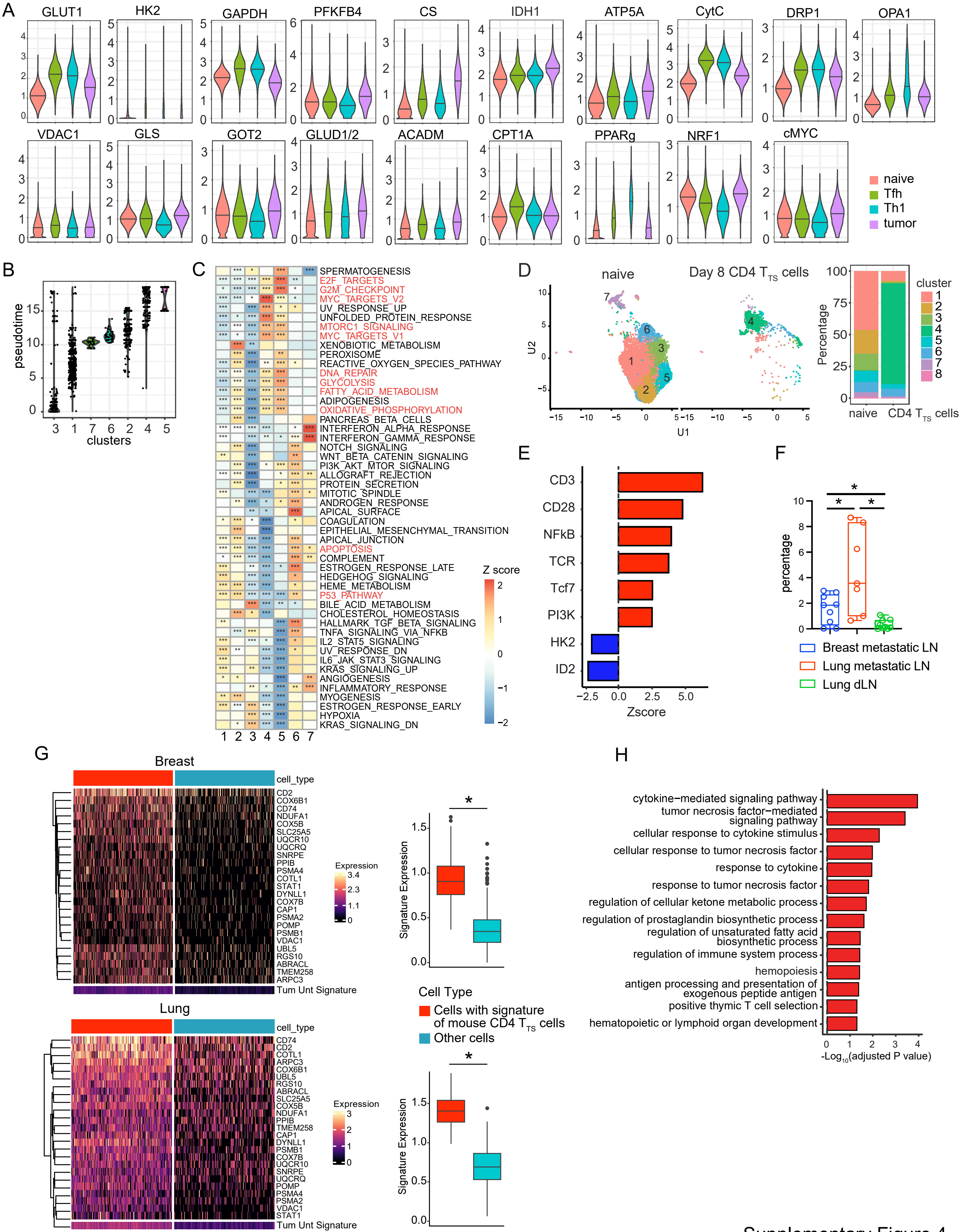


Supplementary Figure 4

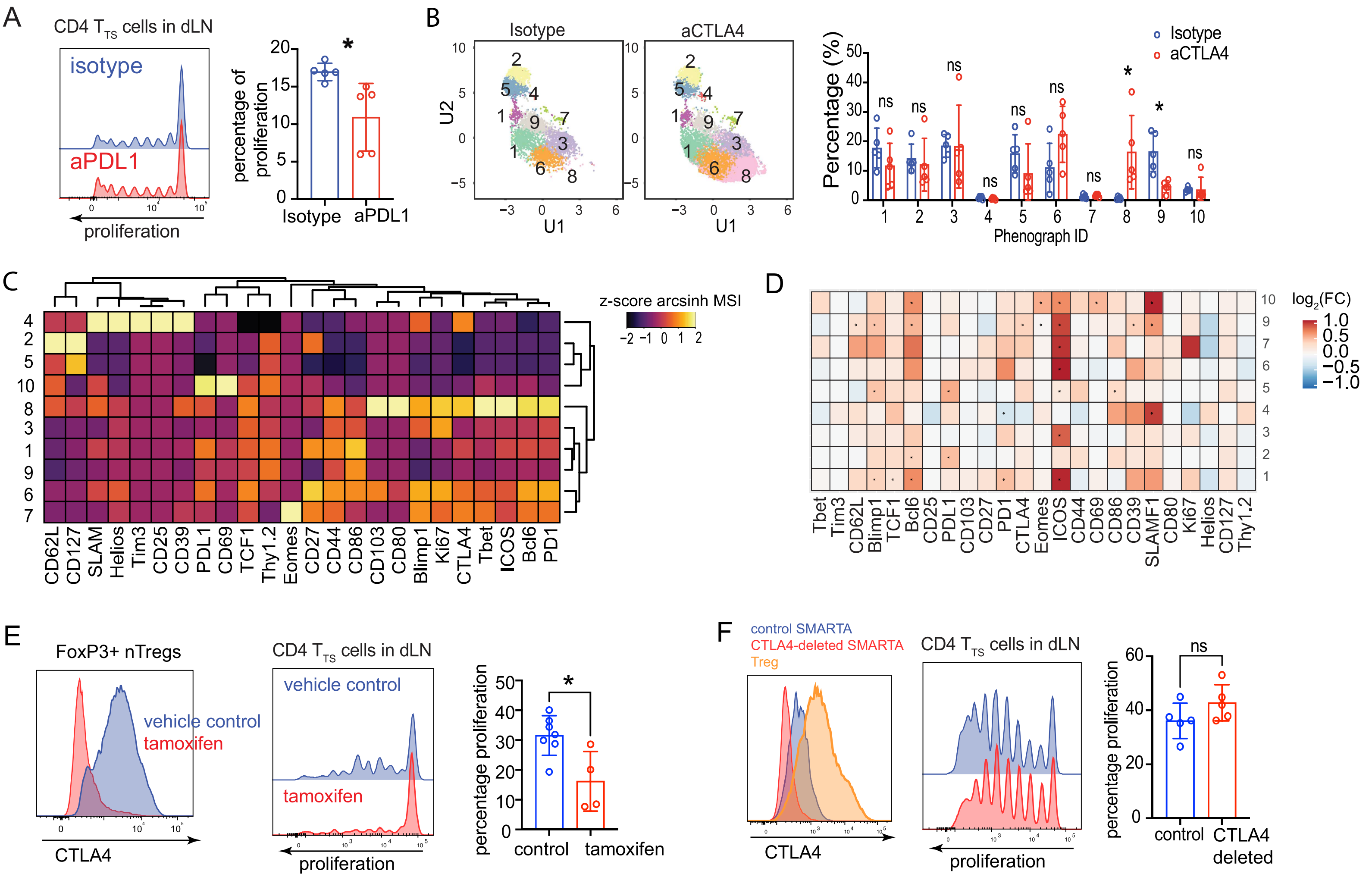
A
CD4 T TS cells in dLN
isotype
aPDL1
proliferation
percentage of proliferation
Isotype
aPDL1
B
Isotype
aCTLA4
U1
U2
Percentage (%)
Phenograph ID
Isotype
aCTLA4
ns
C
z-score arcsinh MSI
CD62L
CD127
SLAM
Helios
Tim3
CD25
CD39
PDL1
CD69
TCF1
Thy1.2
Eomes
CD27
CD44
CD86
CD103
CD80
Blimp1
Ki67
CTLA4
Tbet
ICOS
Bcl6
PD1
D
log2(FC)
Tbet
Tim3
CD62L
Blimp1
TCF1
Bcl6
CD25
PDL1
CD103
CD27
PD1
CTLA4
Eomes
ICOS
CD44
CD69
CD86
CD39
SLAMF1
CD80
Ki67
Helios
CD127
Thy1.2
E
FoxP3+ nTregs
vehicle control
tamoxifen
CTLA4
CD4 T TS cells in dLN
vehicle control
tamoxifen
proliferation
percentage proliferation
control
tamoxifen
F
control SMARTA
CTLA4-deleted SMARTA
Treg
CTLA4
CD4 T TS cells in dLN
proliferation
percentage proliferation
ns
control
CTLA4 deleted

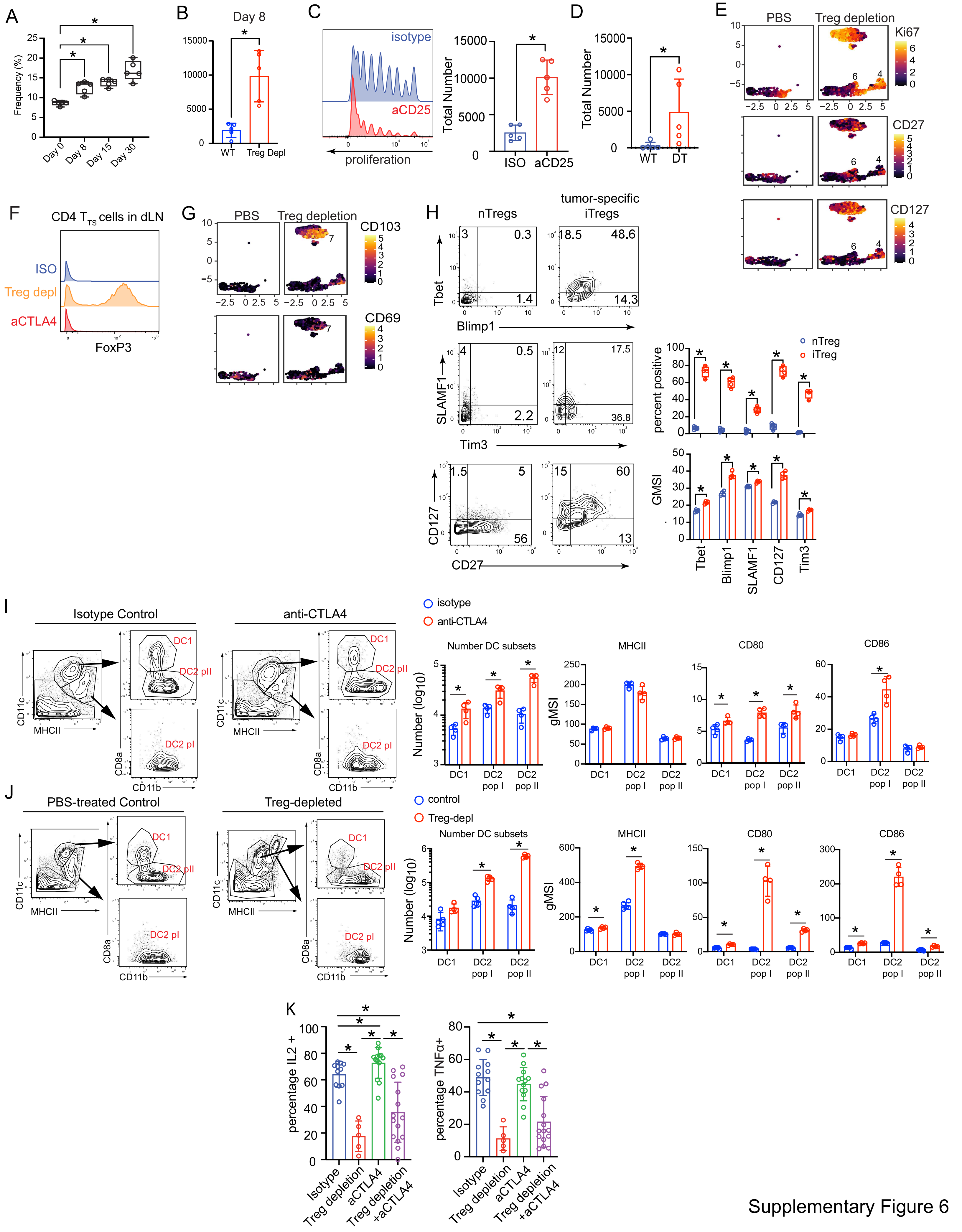


Supplementary Figure 6

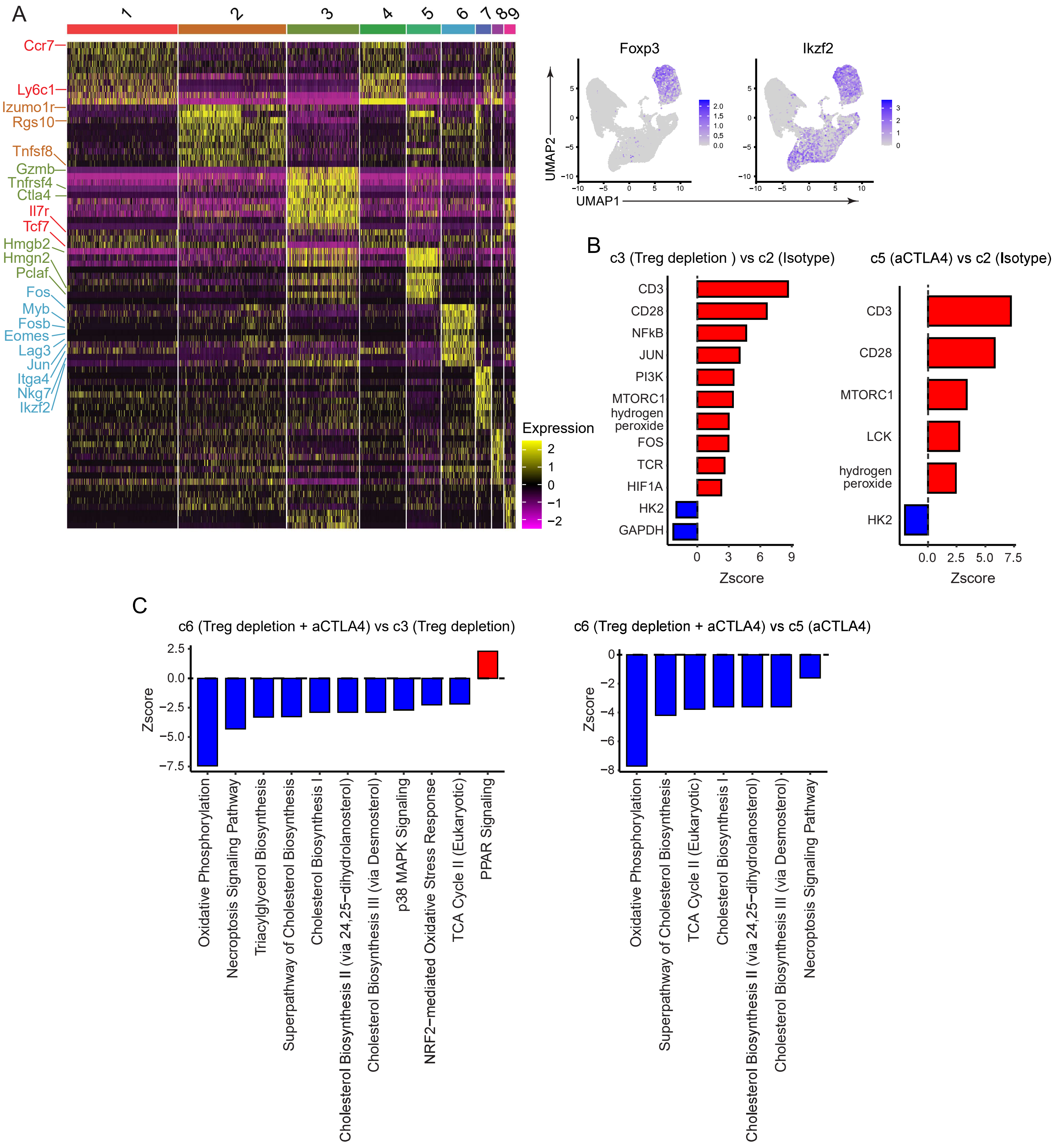


Supplementary Figure 7

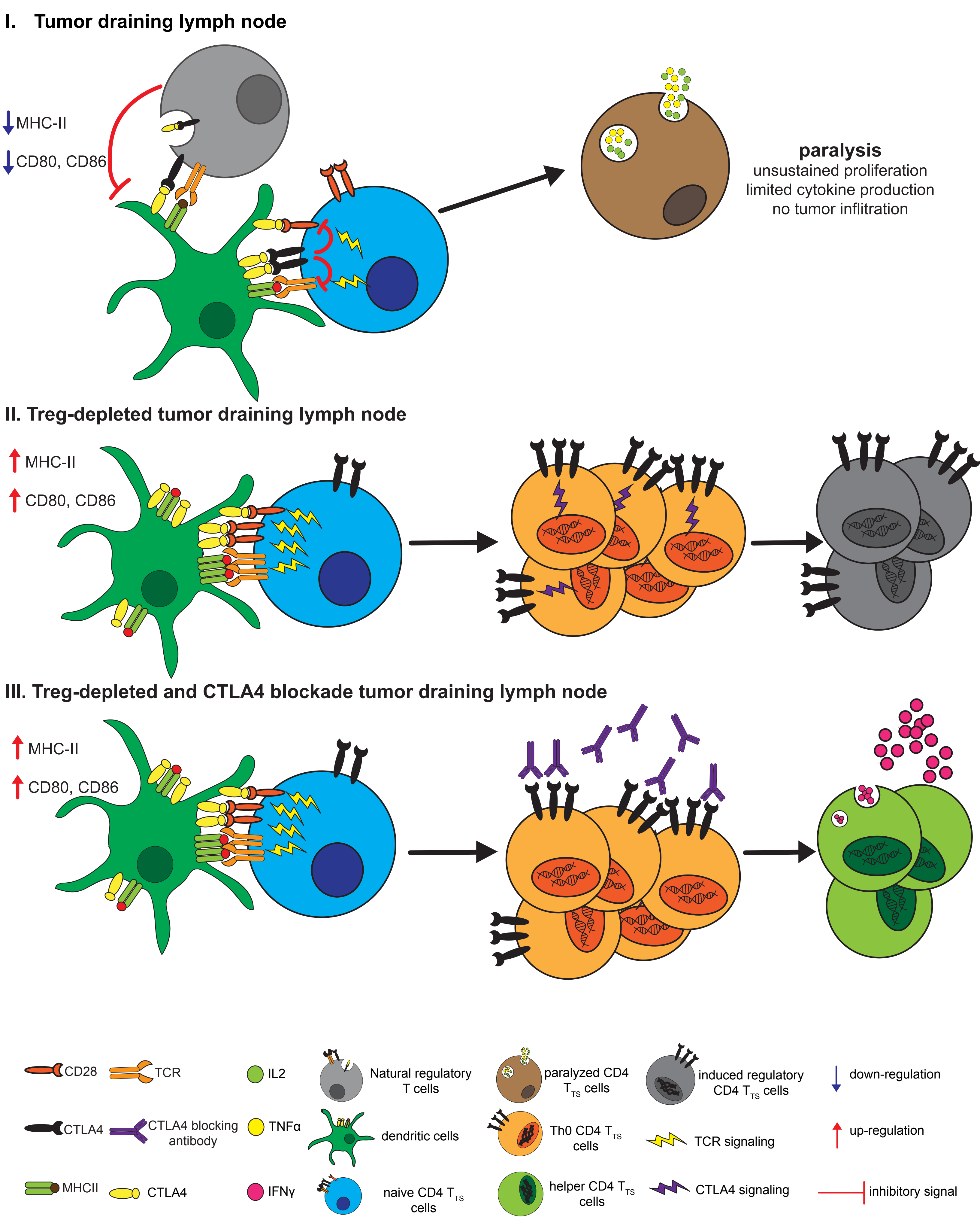

I. Tumor draining lymph node
MHC-II
CD80, CD86
paralysis
unsustained proliferation
limited cytokine production
no tumor inflitration
II. Treg-depleted tumor draining lymph node
MHC-II
CD80, CD86
III. Treg-depleted and CTLA4 blockade tumor draining lymph node
MHC-II
CD80, CD86
CD28
TCR
IL2
Natural regulatory T cells
paralyzed CD4 $T_{TS}$ cells
induced regulatory CD4 $T_{TS}$ cells
down-regulation
CTLA4
CTLA4 blocking antibody
TNFα
dendritic cells
Th0 CD4 $T_{TS}$ cells
TCR signaling
up-regulation
MHCII
CTLA4
IFNγ
naive CD4 $T_{TS}$ cells
helper CD4 $T_{TS}$ cells
CTLA4 signaling
inhibitory signal
